# Development of evolutionarily conserved viral integration sites as safe harbors for human gene therapy

**DOI:** 10.1101/2023.09.08.556857

**Authors:** Marco A. Quezada-Ramírez, Shannon Loncar, Matthew A. Campbell, Krishna M. Parsi, Robert J. Gifford, Robert M. Kotin

**Author notes:** School of Life and Environmental Sciences, The University of Sydney, 380 Werombi Road, Camden NSW 2570, AUS.

## Abstract

Gene transfer into CD34^+^ hematopoietic stem and progenitor cells (HSPCs) involving integrating viral vectors has unpredictable outcomes including potential adverse events like leukemogenesis, resulting from insertional mutagenesis. Therefore, identifying and characterizing genome safe harbor (GSH) sites where exogenous gene material can be safely integrated into adult progenitor and stem cells is critically important for therapeutic gene addition. Here, we present a novel approach to identify new GSH sites based on a proven system of stable transgene insertion: the evolutionarily conserved integration of parvovirus DNA into the germlines of host species. From a dataset of 199 unique endogenous parvovirus (EPV) integration events identified in host species genomes, 102 loci were mapped to the human genome with 17 being experimentally evaluated as GSHs in primary human CD34^+^ HSPCs. Nine promising GSHs resulted in cells edited using nucleofection alone or in combination with rAAV transduction. Of the nine GSH sites, six edited loci displayed sustained transgene expression in both erythroid and immune phenotypes while three clearly exhibited immune branch specific-regulation. Following this approach, additional GSH sites are likely to emerge from the remaining mapped loci for gene addition in hematopoietic stem and progenitor cells. Since it is unlikely that the GSH-lineage-restricted transgene expression is exclusive to hematopoietic stem cells, this approach extends the options for gene knock-ins while reducing the risks of insertional mutagenesis, unpredictable expression profiles, effects on differentiation, and increasing therapeutic effects.

## Introduction

Genome safe harbors (GSH) may be defined at a high-level, as sites within the human genome where new genetic information can be introduced without disrupting normal cellular functions. In contrast, gene therapy applications that utilize integrating viral vectors, e.g., lentivirus or gammaretrovirus, tend to generate insertions randomly into transcriptionally more active and accessible euchromatin regions. Indiscriminate integration results in heterogeneous phenotypes of the transduced cell population with recombination events that can result in genotoxicity, gene silencing, unpredictable transgene expression or stability, loss of ‘“stemness”, or dysregulated cellular proliferation, i.e., leukemias ^1–3^. Thus, identifying GSHs that provide reliable locations for gene addition becomes a high priority for gene and cellular therapies. Clinical outcomes of lentiviral vector-dependent gene therapies addressing blood disorders (e.g. multiple myeloma ^4^, β-hemoglobinopathies^5^, coagulopathies^6^, Fanconi anemia^7^, etc.) would improve upon selection of an appropriate GSH from an available catalog of verified GSH sites that reduce the risks associated with cell engineering. Importantly, the accessibility to primary human CD34^+^ hematopoietic stem and progenitor cells (HSPCs) combined with programmable endonuclease technologies (particularly CRISPR/Cas9) constitute clinically relevant model to assess candidate GSHs for autologous cell therapies.

To date, efforts are ongoing to find new human GSH sites beyond the widely applied *AAVS1*^8–10^, *CCR5*^11,12^, and *hRosa26*^13–15^ loci. Criteria have been proposed for selecting potential GSH sites that are intergenic and preferentially 50-300 kb distant from known or predicted transcriptional units^16–19^. However, the number of characterized sites following this approach have been limited and mostly evaluated in established cell lines with few examples using primary human cells^20,21^. Other criteria proposed for identifying GSH sites are based on using endogenous mobile elements^22^ or higher-order, chromatin architecture analysis^21,23^ but outcomes have not been particularly effective. Thus, *AAVS1* which was developed as a GSH ^10,15^, remains a popular and widely used locus for directed transgenesis given that its knock-in does not affect the cell viability or differentiation ^24,25^ although some reports argue that transgene silencing occurs in a cell lineage^26^ or promoter-specific manner^27^. The *AAVS1* site was originally identified as an AAV provirus integration site in exon 1 of *PPP1R12C* gene on human chromosome 19, within a motif consisting of AAV Rep protein-dependent origin of DNA synthesis ^8,9,28^, however, commonly available editing reagents relocated the site to the first intron of *PPP1R12C*^29^. As *AAVS1* is a parvovirus integration site, we hypothesized that other parvoviral integration loci may predict GSH sites too.

Parvoviruses are among the small number of viruses known to have contributed genes to the vertebrate germline via viral integration and horizontal gene transfer^30^. As a result, endogenous parvoviral elements (EPVs) have resulted from ancient infections across phylogenetically diverse host species that integrated into the germline and persisted during multiple speciation events across geologic time scale^31^. An extensive phylogenetic analysis of EPVs in vertebrate genomes disclose clear homology to members of extant parvovirus genera, including *Amdoparvovirus* ^32^, *Protoparvovirus* ^33,34^, and *Dependoparvovirus* ^31,35^. The viral DNA endogenization represents an alternative to random mutation where large-step mutations resulted in the acquisition of novel viral “alleles,” facilitated genomic rearrangements, disrupted gene expression, created novel transcription factor binding sites, or resulted in “exaptation” of viral genes with potential benefits to the host organisms^36^, notably syncytin - a retrovirus envelope protein co-opted by placental mammals for trophoblast fusion early in embryogenesis^37^. Thus, identifying human orthologous sites to EPVs in non-human vertebrate genomes may indicate suitable sites for gene addition.

In this study, we identify 102 potential human GSH sites that are orthologs to EPV integration sites in the respective host species^38^. Orthologous loci were broadly categorized as intergenic or intronic according to the relative location in the host species and in the human genome. Additionally, the accessibility for genome editing in hematopoietic precursors was predicted by analyzing available ATAC-seq data. Through CRISPR/Cas9-mediated genome editing, seventeen of these loci (8 intergenic and 9 intronic) were evaluated as GSH sites by inserting an *eGFP* expression cassette in primary human CD34^+^ HSPCs. The edited cells displayed no alterations in self-renewal or multipotency, i.e., stemness, demonstrating suitability of these loci for gene addition. Moreover, transgene expression was maintained across differentiation stages, indicating stable integration. Remarkably some GSHs displayed immune cell restrictive regulation. To our knowledge, this is the first study that provides an extended catalog of human GSH sites with potential applications for CD34^+^ HSPC-based therapies. Furthermore, it is likely that lineage restrictive expression is not exclusive to HSPCs and can be extended to other tissues with additional GSH sites that are likely to emerge from the remaining mapped loci.

## Results

### Homology ascertainment relative to human genome

The local genomic landscape surrounding EPV insertions may be broadly categorized as intergenic or intronic. This categorization carries over when identifying the equivalent orthologous position in the human genome. For the 102 mammalian EPV loci (62 intergenic and 40 intronic) we mapped the orthologous locus in the human genome by comparative genomics, regional genome alignments and gene synteny^39^, as well as similarity of DNA flanking EPVs using BLAT or BLASTN analysis. Relative positions between host species and human genome were established by interspecific collinearity and were refined to more precise coordinates based on sequence homology.

The putative GSH loci identified via this approach were found distributed across the human genome without obvious sequence or higher order common features (Fig. 1a; Supplementary Table 1). The majority of the EPVs were derived from members of the *Dependoparvovirus* and *Protoparvovirus* genera (Fig. 1b). An EPV loci nomenclature referencing the parvoviral genus from which each EPV is derived was established and can be consulted together with coordinates of their relative position on the human genome in the Supplementary Tables 1 and 2.

**Fig. 1.**
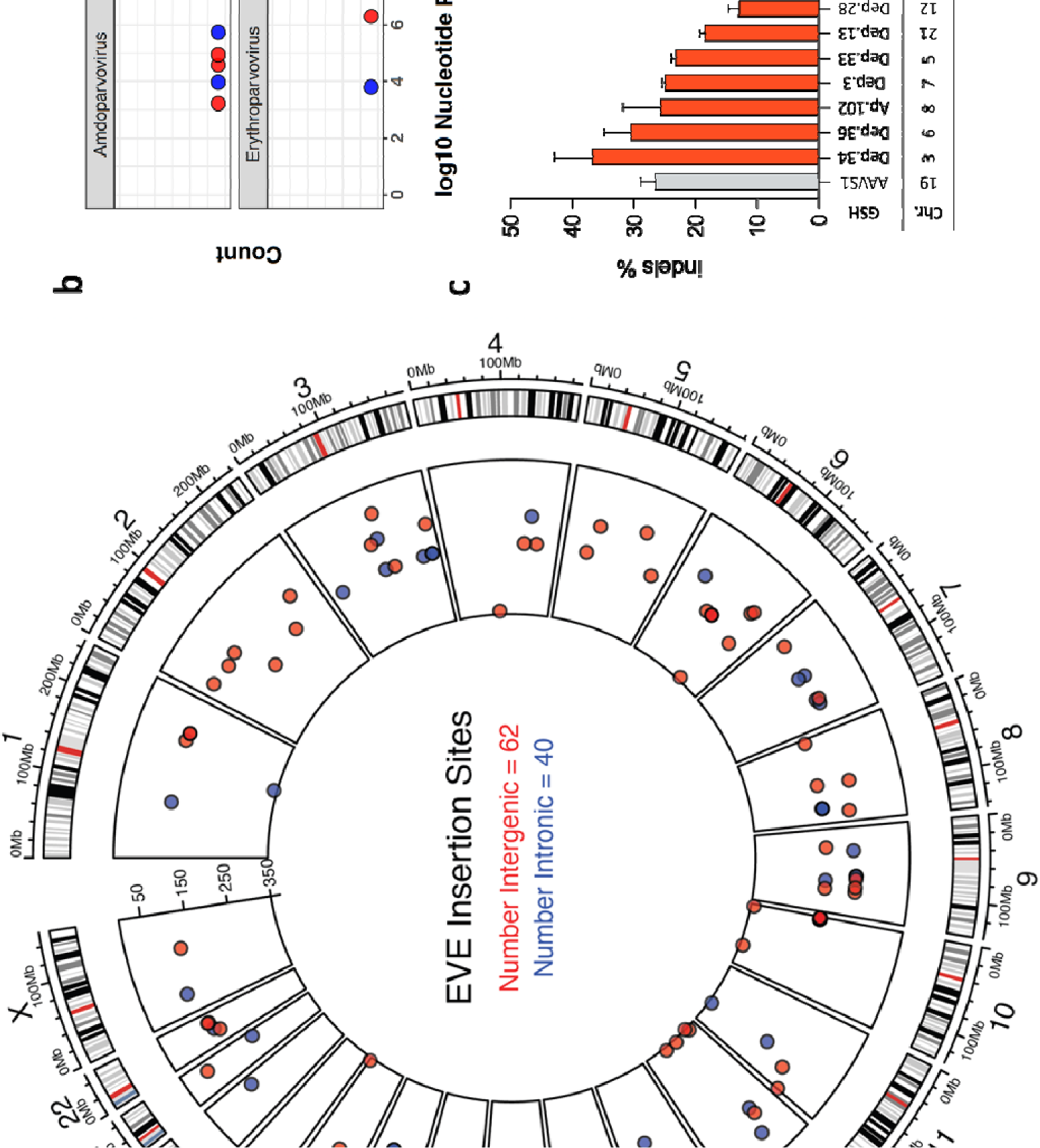
Identification of 102 potential genome safe harbor (GSH) sites in the human genome. **a.** Circle plot of EPV orthologues mapped onto the human genome. The outer layer represents human chromosomes, 1 to 22 and X. Chromosome lengths are indicated by the tick marks, e.g., 0 - 100 Mb. Black and red radial marks represent chromosomal banding patterns and centromeres, respectively. The inner layer depicts intergenic and intronic locations of GSHs as red and blue circles, respectively, with the estimated divergence time of the host species taxa from humans scaled from 0 to 350 million years ago (MYA). **b.** Comparative genomic analysis of the precision of homology between EPV host species and human genome. EPVs are derived from four parvovirus genera: *Amdoparvovirus*, *Dependoparvovirus*, *Erythroparvovirus* and *Protoparvovirus*. The distribution displays abundance of *Dependoparvovirus* and *Protoparvovirus* related loci. **c.** Propensity to genome editing of 17 selected human EPV orthologs evaluated by indel formation in HEK293T cells following the expression of *Cas9* and a specific gRNA. The intergenic or intronic category is depicted as red and blue bars, respectively. The nomenclature of the assessed GSH sites (abbreviated EPV name) used for this report as well as the chromosomal host are indicated on the x-axis. Indel frequencies were determined through TIDE decomposition analysis. Mean ± s.d., *n* = 2 independent experiments.

### Selection of candidate GSH sites

Human ortholog loci from the intronic and intergenic categories were selected for experimental characterization (Supplementary Table 2). The proximity to open chromatin regions was evaluated using available ATAC-seq data for human CD34^+^ HSPCs^40^, with sites within 2 kb from an ATAC-seq peak prioritized for evaluation. Seventeen loci, 8 intergenic and 9 intronic, were selected for the first round of screening in HEK293T cells (Supplementary Table 2). Initially, single guide RNAs (hereafter gRNAs) were designed using online gRNA predictive tools (CHOPCHOP, CRISPOR, and CRISTA). Then, three to five highly scored gRNAs per locus (with minimal off-target prediction) were selected and incorporated into pX330 plasmids to co-express SpCas9 and the corresponding gRNA. HEK293T cells were independently transfected to select the best gRNA per locus using a highly cited *AAVS1* gRNA for comparison purposes^41^ (Fig. 1c, Supplementary Table 3). Relative to *AAVS1* indel formation the GSH candidate loci indel frequencies suggested reasonable propensity to genome editing (Fig. 1c), therefore, we proceeded with a proof-of-concept study editing primary human CD34^+^ HSPCs derived from bone marrow of healthy adult donors.

### Editing of GSH candidates in human CD34^+^ HSPCs with CRISPR/Cas9 and ssDNA templates

Several conformers of DNA templates have been reported for targeted integration into hematopoietic cells. These include reporter or therapeutic genes in plasmids or linear double stranded DNA (dsDNA) flanked by homology arms that mediate the homologous recombination (HR) ^20,42^, or flanked with *gRNA* recognition sites for homology-independent targeted integration (HITI) ^43^. However, these dsDNA templates are transcriptionally competent as episomal DNA and elicit robust innate immune responses that may result in cytotoxicity. On the other hand, recombinant AAV6 vectors have been developed for HR and HITI but even when these vectors enhance the genome editing outcomes ^44^, the DNA tends to remain episomal after the transduction and positive or negative strands that may anneal into transcriptionally active duplex DNA potentially leading to false positive editing results. Therefore, we decided to test single-stranded DNA (ssDNA) templates as these reportedly correct point mutations associated with blood disorders ^45,46^. Our preliminary comparison of dsDNA and ssDNA templates displayed reduced GFP background with ssDNA templates (Supplementary Fig. 1A and B), reducing the random integration effects of episomal DNA templates following transfection (Supplementary Fig. 1C and D). Consequently, we engineered ssDNA templates per locus composed of an expression cassette consisting of *eGFP* open reading frame regulated by the MND synthetic promoter and rabbit beta-globin polyadenylation signal. The expression cassette was flanked by locus specific 300 nt homology arms yielding approximately 2 kb templates (Fig. 2a).

**Fig. 2.**
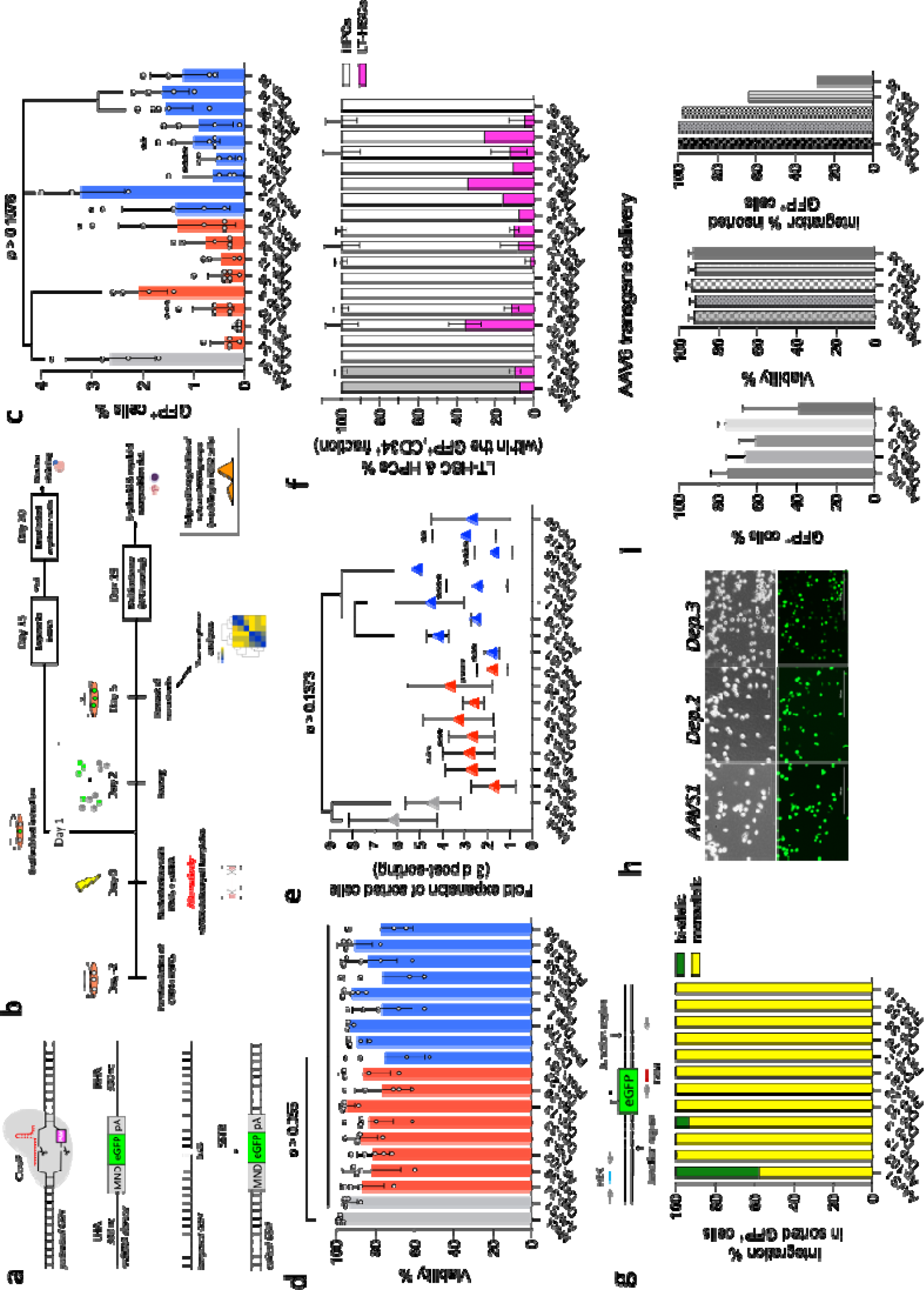
Targeting of GSH candidates in primary human CD34^+^ HSPCs. **a** Genome targeting with *eGFP* expression cassettes into GSH candidates using locus specific CRISPR/Cas9-ssDNA editing sets. Single-stranded DNA donor templates were flanked with 300 nt homology arms to mediate homologous recombination through single stranded template repair (SSTR) pathway. **b** Experiments with bone marrow CD34^+^ HSPCs. Pre-stimulated cells were nucleofected in presence of the CRISPR/Cas9-ssDNA editing set and cultured for two days before the sorting of the GFP^+^ population. The GFP^+^ cells were cultured for an additional three days before downstream processing. Alternatively, AAV6 vectors were used to deliver templates to target *Ap.102*, *Dep.33*, *Dep.13*, *Dep.1* and *Dep.35* loci. In parallel, 1 day-nucleofected cells were cultured in erythroblast expansion conditions before maturation with human EPO. **c** Percentage of GFP+ cells at two days post-nucleofection. Intergenic and intronic loci are depicted as red and blue bars, respectively. Open circles represent individual human donors. *AAVS1* GSH (gray) served as reference control. Mean ± s.d, ANOVA followed by Dunnett test * *P* < 0.05, ** *P* < 0.01, *** *P* < 0.001 and **** *P* < 0.0001. *n* ≥ 4 independent experiments. **d** Cell viability determined by flow analysis of propidium iodide stained cells. Comparison control corresponds to unedited cells (mock). Open circles represent independent human donors. Mean ± s.d, ANOVA followed by Dunnette test, *n* ≥ 3 independent experiments. **e** Proliferation rate of GFP^+^ HSPCs three days after sorting. ANOVA followed by Dunnett test, *n* ≥ 3 independent experiments. **F** Targeting of long term-HSCs (CD34^+^/CD90^+^, magenta) and progenitor cells (CD34^+^/CD38^+^, CD34^+^/CD38^-^) within the GFP^+^/CD34^+^ fraction. n = 2-3 independent experiments using different human donors. **g** Targeted Integration in GFP+ cells sorted after 14 days of liquid culture. Specific primer/probe sets (arrows and bars) were utilized. The efficiency was calculated as the FAM/HEX ratio. *n* = 2 independent experiments. **h** Persistent transgene expression after 45 days of liquid culture. **i** AAV6 delivering of templates for *Ap.102*, *Dep.33*, *Dep.13*, *Dep.1* or *Dep.35* loci show increased levels of sorted GFP^+^ cells without affecting cell viability and high level of integration. Mean ± s.d., *n* = 2 independent experiments.

Thus, synthetic gRNAs complexed with SpCas9 were co-delivered together with 3 µg of the corresponding ssDNA template into 2.5×10^5^ primary human CD34^+^ HSPCs following a nucleofection protocol (Lonza Inc., Morristown, NJ, USA). To minimize transcriptional interference, intronic GSH loci templates were engineered to insert the transgene in opposite orientation relative to the transcriptional unit hosting the candidate GSH site. Two days after nucleofection, the GFP^+^ cells were collected through FACS sorting and cultured in SFEM liquid medium for additional 3 days (Fig. 2b). Unless noted differently, the experiments with undifferentiated CD34^+^ HSPCs spanned seven days to avoid impairing the stemness features that occur following prolonged cell culture ^47^. The GFP^+^ percentages ranged from 0.12% ± 0.05 GFP^+^ cells (mean ± s.d. *P* < 0.0001), for the intergenic locus *Dep.36*, up to 3.23 % ± 0.86 GFP^+^ cells (*P* = 0.8938) for the intronic locus *Dep.2* (Fig. 2c). The reference control for these experiments, locus *AAVS1*, reached 2.65% ± 0.88 GFP^+^ cells (Fig. 2c). Not surprisingly, the GFP^+^ percentages in CD34^+^ HSPCs differed substantially from the values estimated through indels in human embryonic kidney (HEK) 293T cells that were typically ≥ 10% (Fig. 1c). Multiple factors account for the different outcomes including the HEK293T-suppressed innate immune responses, overall robustness of this cell line, and hetero-/euchromatization differences between cell types. For instance, the candidate GSH site *Dep.36* (intergenic) displayed high propensity to genome editing in HEK293T with approximately 30% indels formed (Fig. 1c), but yielded the lowest percentage of GFP^+^ HSPCs after the nucleofection (Fig. 2c). Although we consulted available ATAC-seq data for CD34^+^ HSPCs^40^ as an additional criterion to select the best GSH candidates, no evident correlation was observed between editing efficiencies and the proximity to reported open chromatin spots (Supplementary Tables 2 and 3). An extreme example, is the *Dep.2* intronic locus that resulted in the highest percentage of GFP^+^ cells at two days post-nucleofection (3.23% ± 0.86, *P* = 0.8938) despite having 15 kb distance from the nearest ATAC peak (Supplementary Table 3). Furthermore, persistent *eGFP* expression was monitored through differentiated stages of edited *Dep.2* cells, a phenomenon replicated by eight other loci lacking proximity to reported ATACseq peaks, namely the intergenic loci *Dep.3*, *Dep.22*, *Ap.102*, and the intronic sites *Dep.55*, *Prot.218*, *Prot.2*, *Prot.181* and *Dep.1* (Supplementary Table 3; Fig.2c and Fig. 4a).

The cell viability fluctuated between 75.15% ± 20 live cells (mean ± s.d., *P* = 0.0551), for the targeted *Prot.181* site, and 94.65% ± 3.71 live cells (*P* = 0.9994) for the targeted *Dep.13* site, as compared to the 98.4% ± 0.92 of the mock control (i.e., electroporated cells in absence of any CRISPR/Cas9-ssDNA editing set) (Fig. 2d). The proliferation rate decreased significantly within the three days following the sorting of GFP^+^ HSPCs compared to mock control, 6.19-fold ± 1.96, excepting for *AAVS1* (4.43-fold ± 1.22, *P* = 0.3294), *Dep.1* (4.56-fold ± 1.53, *P* = 0.4331), *Dep.2* (4.21-fold ± 0.50, *P* = 0.1373), and *Dep.35* groups (5.17-fold ± 0.18, *P* = 0.9885) (Fig. 2e). Despite this, large colonies of differentiated cells formed independently of the edited GSH locus demonstrating that neither genome modification nor *eGFP* transgene expression impaired the proliferation capacity of the cells (Fig. 4a).

### Targeting Long Term-HSCs CD34^+^ populations

Primary CD34^+^ HSPCs are a heterogeneous cell population whose developmental stages are identifiable by specific cell-surface markers. Within this, the long-term hematopoietic stem cells (LT-HSCs) are those with self-renewal and long term engraftment potential considered the most clinically relevant for autologous therapies ^48^. In bone marrow, HSPCs are very scarce, estimated at 0.01% the cell population and LT-HSC in bone marrow has been estimated to be even scarcer, approximately 0.0001% of the cell population. To determine whether GSHs were targeted in LT-HSCs and cells survived gene addition, we analyzed the GFP^+^ cell fraction to distinguish potential LT-HSCs (CD34^+^/CD90^+^) from the most abundant progenitor cells (CD34^+^/CD90^-^/CD38^+^, CD34^+^/CD90^-^/CD38^-^) ^49^. Interestingly, the results suggest that not all loci are accessible in potential LT-HSPC populations (Fig. 2f). The loci that produced edited LT-HSPCs include *AAVS1* (10.32%), *Ap.102* (35.97%), *Dep.3* (12.07%), *Dep.28* (2.05%), *Dep.22* (8.56%), *Prot.181* (10.59%), *Dep.2* (8.33%), *Prot.176* (16.67%), *Dep.1* (34.7%), *Prot.2* (11.18%), *Dep.35* (13.22%), *Dep.55* (26.09%), and *Prot.218* (5.6%) (Fig. 2f), making these attractive candidates for long term animal studies.

### GSH targeted integration in human CD34^+^ HSPCs

To verify template integration into the GSH candidate loci, genomic DNA was extracted from GFP^+^ HSPCs sorted after 14 days of liquid cell culture. The DNA was analyzed by droplet digital PCR (ddPCR) with a locus-specific primer/probe set spanning the junction region and a second primer/probe set specific for an unedited part of the locus to obtain the total number of alleles (edited and unedited) in the sample (Fig. 2g; Supplementary Fig. 2; Supplementary Table 7). Considering two GSH alleles per diploid genome, a 50% ddPCR signal represents one hundred percent of monoallelic integration whereas >50% is indicative of partial bi-allelic integration. Thus, the results indicate that the editing of *AAVS1* (reference locus) occurred with 40% of bi-allelic integration (Fig. 2g), whereas the targeting of intergenic (*Dep.3*, *Dep.13*, *Dep.33*, and *Ap.102*) and intronic candidates (*Dep.1*, *Dep.2*, *Dep.35*, *Prot.2*, *Prot.181*, *Prot.176*, and *Prot.218*) resulted mostly in monoallelic integration likely resulting from more restricted accessibility (Fig. 2g). Notwithstanding, the integration of the transgene was stable for at least 45 days as demonstrated by the long-term culture of *AAVS1*, *Dep.2* (intronic), and *Dep.3* (intergenic) edited cells (Fig 2h). On the other hand, the relatively low level of GFP^+^ cells obtained from the targeting of *Dep.34*, *Dep.36*, *Dep.28*, and *Dep.56* loci (Fig. 1c) resulted in difficult and sometimes undetectable integration events. Factors accounting for this outcome might include inaccessibility to the editing components, unfavorable ssDNA template secondary structure (e.g. the donor DNA template for the intronic *Dep.56* site displayed energetically stable secondary conformations), disruption of vital pathways, or even HUSH-mediated repression of intronless transgenes ^50^, making them poor candidates for future *in vivo* transplantation studies.

Tracking studies of corrected HSPCs in Wiscott-Aldrich syndrome or β-hemoglobinopathy patients have estimated that as few as 98 corrected HSPCs per 10^6^ infused CD34^+^ cells are able to engraft the bone marrow for long term-production of granulocytes^48^. Given that our experiments were designed using relatively small cell numbers (2.5×10^5^ to 1×10^6^ CD34^+^ HSPCs per single nucleofection targeting one candidate GSH site), the results suggest that the scaling-up to clinical preparation numbers (usually >5X10^6^ electroporated cells) would increase the bioavailability of therapeutic cells ^44,51^. Alternatively, if low cell numbers are involved, the rAAV6 vectors reportedly substantially increase editing efficiencies. To test the latter approach, we combined nucleofection and rAAV6 transduction to deliver homology arm-carrying templates targeting three intergenic (*Ap.102*, *Dep.33* and *Dep.13*), and two intronic GSH loci (*Dep1* and *Dep.35*) which showed low efficiencies following ssDNA template nucleofection (Fig. 2c and Fig. 2i). Equivalent cell numbers were used (i.e., 2.5×10^5^ cells). As expected, at two days post transduction, the percentage of sorted GFP^+^ cells increased substantially without affecting cell viability (Fig. 2i), but given that AAV transgenes persist episomal after internalization, accurately determining the integration level on sorted cells requires further analysis (Fig, 2h and Supplementary Fig. 1E). Regardless, these results demonstrate the feasibility to increase the targeting of apparently difficult GSH candidate loci.

### Transcriptional changes in edited CD34^+^ HSPCs

To determine whether and to what extent GSH editing perturbs the transcriptional homeostasis of stem and progenitor cells, RNAseq was performed with the bulk of sorted GFP^+^ HSPCs from three intronic (*Dep.2*, *Dep.55* and *Prot.218*) and two intergenic (*Dep.3* and *Ap.102*) GSH loci targeted with ssDNA templates. Differentially expressed genes (DEGs), were determined by comparing edited versus unedited control cells from the same human donor (mock nucleofection). Consequently, the targeting of the reference GSH site, *AAVS1,* resulted in 1,595 DEGs, whereas the intronic GSH candidates *Dep.2*, *Dep.55*, and *Prot.218*, resulted in 2,163, 925 and 1,144 DEGs, respectively (Figure 3a and 3b). Perhaps unsurprisingly, the genome editing of the intergenic GSH sites *Dep.3* and *Ap.102* resulted in fewer DEGs reporting 509 and 496, respectively (Fig. 3a and 3b). Thus, even though the CD34^+^ HSPCs constitute a heterogeneous population the results suggest that targeting intergenic GSH sites tends to perturb the transcriptome to a lesser extent than intronic GSH loci.

**Fig. 3.**
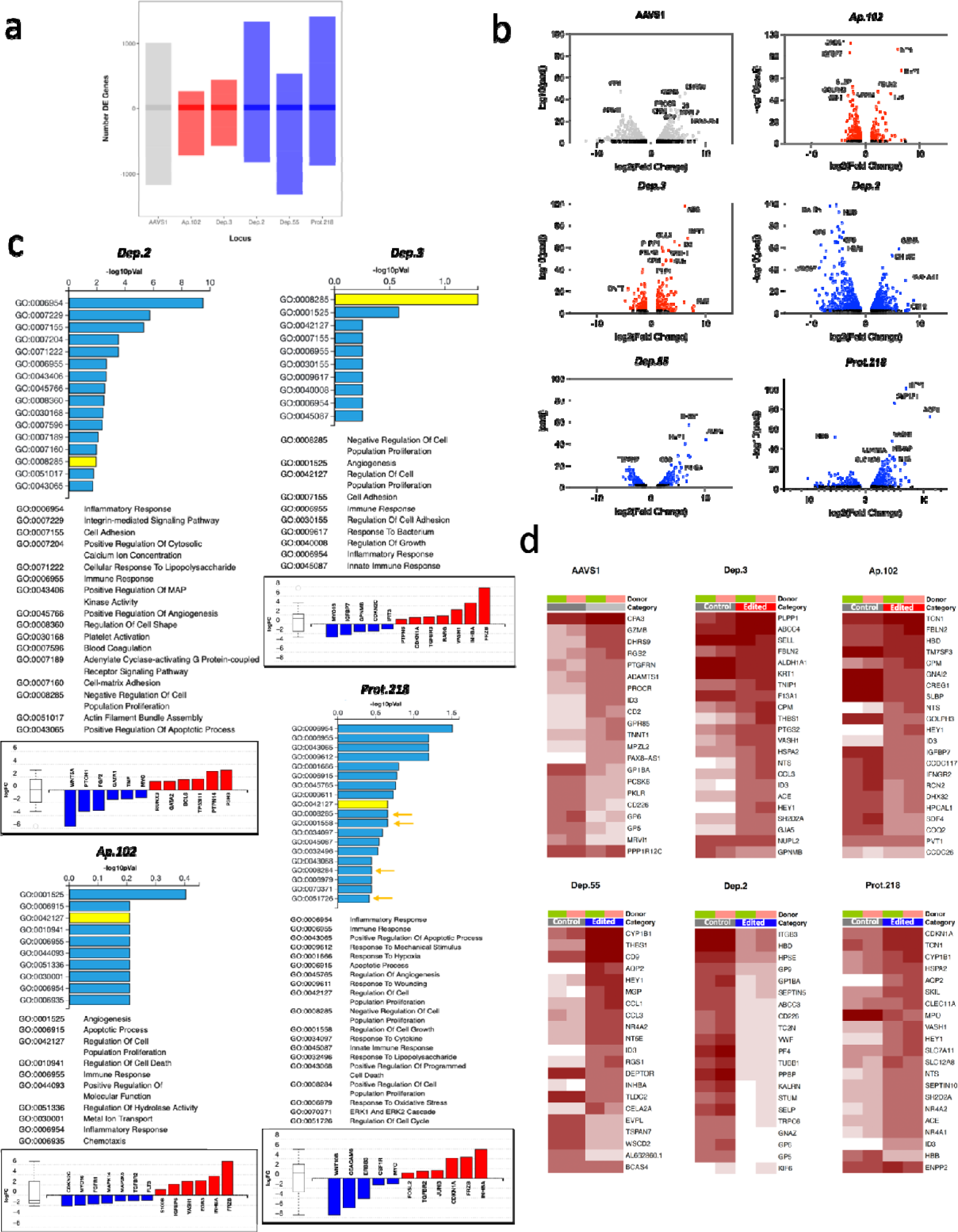
Gene expression changes following the targeting of GSH candidates in primary human HSPCs. **a** Comparison of total differentially expressed genes (DEGs) among intergenic (*Dep.3*, *Ap.102*) and intronic (*Dep.2*, *Dep.55*, *Prot.218*) GSH candidates, red and blue symbols, respectively, compared to the reference GSH, *AAVS1,* shown in gray. Genes consistently upregulated (n = 30) and downregulated (n = 15) across all experiments are indicated by shading in barplots. **b** Volcano plots of the global transcriptional changes after genome editing of *AAVS1* (1008 up-/587 downregulated genes), *Ap.102* (133 up-/363 downregulated genes), *Dep.3* (220 up-/289 downregulated genes), *Dep.2* (1342 up-/821 downregulated genes), *Dep.55* (269 up-/656 downregulated genes) and *Prot.218* (798 up-/436 downregulated genes). Genes with adjusted *p*-values < 0.01 are highlighted in color, *AAVS1* gray, intergenic red and intronic blue. Highly significant DEGs are named. **c** Gene ontology annotations for DEGs obtained from *Dep.2*, *Dep.3*, *Ap.102*, and *Prot.218* editing. Genes with attributes to regulation of cell proliferation are indicated by yellow bars with representative genes plotted at the bottom, blue downregulated oncogenes, red upregulated tumor suppressors. **d** Heatmaps with the top 20 more significant DEGs per edited locus (all DEGs are available in a database uploaded on NCBI). The host gene for intronic loci or the flanking genes for intergenic loci are appended at the bottom. Color scale represents the Log_2_ of normalized counts of *n* = 2 independent donors (pink and green boxes).

In addition, no substantial transcriptional changes on the genes proximal to the insertion sites were detected (Supplementary Fig. 3). Uniquely, the host gene for the intronic *Prot.218* locus (*ENPP2* gene) displayed a modest 2.95-fold increase on gene expression (*P* = 2.4×10^-5^) whereas a downstream transcript (*DEPTOR*) decreased by 0.18-fold (*P* = 2×10^-11^) (Fig. 3d and Supplementary Fig. 3). Similarly, transcripts mapping to *GPNMB*, downstream to *Dep.3* locus, displayed a fairly robust, 0.34-fold decrease relative to unedited control cells (*P* = 0.03) (Fig. 3d and Supplementary Fig. 3).

The gene ontology (GO) analysis of DEGs ranked more than one group of genes associated to regulation of cell proliferation (GO:0008285, GO:0042127, GO:0001558, GO:0008284, and GO:0051726). Notably, the genes associated with malignant transformation showed no evidence of alteration (Fig. 3c). The evaluated transcriptomes revealed upregulation of tumor suppressor genes *CDKN1A* (8.52-fold change, *P* ≤ 10^-8^), *INHBA* (29.1-fold change, *P* ≤ 0.001), and *TP53I11* (2.5-fold change, *P* ≤ 0.01) (Figs. 3b and c). In contrast, RNAseq analysis showed downregulation of leukemia-associated proto-oncogenes such as *CCND1/2* (0.46-fold change, *P* < 0.05), *CCNE2* (0.38-fold *P* = 0.0076)*, MYC* (0.42-fold change, *P* < 0.001), *SRC* (0.3-fold change, *P* < 10^-5^), or *MPL* (0.25-fold change, *P* < 0.05) (Fig. 3c). Additionally, intergenic GSH sites *Dep.3* and *Ap.102* displayed downregulation of the AML-associated proto-oncogene *FLT3* (0.44-fold change, *P* ≤ 0.0014) and overexpression of the tumor suppressor gene *INHBA* (18.4-fold change, *P* ≤ 0.005) (Fig. 3c). With the exception of the reference *AAVS1* edited cells, all the GSH-targeted cells underwent overexpression of *HEY1* (98.8-fold, *P* < 10^-40^, for *Dep.3*, *Dep.55*, *Ap102*, and *Prot,218*), or upregulation of *HES7* (8.9-fold *P* = 0.017 for *Dep.2*), both transcriptional repressors for the maintenance of blood precursors (Fig 3d). In addition, overexpression of *FRZB* (74.2-fold change, *P* < 0.01), an inhibitor of the □-catenin pathway, was observed (Fig. 3b and 3d). Similarly, all the edited GSH groups displayed up-regulation of genes associated with innate immune response presumably resulting from *in vitro* manipulation, i.e., nucleofection of ssDNA, CRISPR:Cas9 complex, etc. (Fig. 3c). Together these changes confirmed the observed phenotype of the manipulated CD34^+^ HSPCs in culture (i.e., reduced cell proliferation, Fig. 2e) and indicate low propensity for malignant transformation, however, deeper sequencing (with homogeneous stem cell systems) as well as cell transformation assays will help to clarify the cellular response to and assess risks associated with editing of the novel GSH sites.

### Edited CD34^+^ HSPCs retained the multipotency with widespread or lineage-specific transgene expression

To determine whether gene addition and transgene expression affected the HSPC proliferation or differentiation, cells were examined following 14 days of culture in differentiation medium. No morphological abnormalities were detected among the colonies (CFUs) derived from manipulated HSPCs (Fig. 4a). Instead, differentiated edited cells formed colonies that continued to express *eGFP*. *Dep.2*, *Dep.3*, *Prot.218*, *Prot.2* and *Prot.181* edited cells produced GFP^+^ colonies of four lineages, erythroid (BFU-E), granulocyte (CFU-G), granulocyte-macrophage (CFU-GM) and macrophage (CFU-M) as was done for the reference group of edited *AAVS1* cells (Fig. 4a). In contrast, although *Dep.35* and *Proto.176*-derived colonies also showed widespread expression across the four lineages, the GFP^+^ signal was dramatically reduced upon differentiation (Fig. 2g and Fig.4a) indicating downregulation of *eGFP* from these loci. Other edited loci resulted in *eGFP* expression biased towards granulocyte/macrophage lineages based on visual inspection of the colonies (*Dep.13*, *Dep.1*, *Dep.55*, *Ap.102* and *Prot.176*) (Fig.4a). Despite dissimilar expression phenotypes, the total colony distributions were similar among all the groups as compared to unedited control cells (Fig. 4b). Interestingly, the edited GSH cells with higher GFP signal and widespread *eGFP* expression by 14 days of differentiation were the reference locus, *AAVS1*, and *Dep.3*, *Dep.2*, *Dep.55*, and *Prot.218* loci (Fig. 4a and 4c) being included as well among those loci resulting in targeting of LT-HSCs (Fig. 2f). These characteristics are attractive for gene addition in human CD34^+^ HSPCs.

**Fig. 4.**
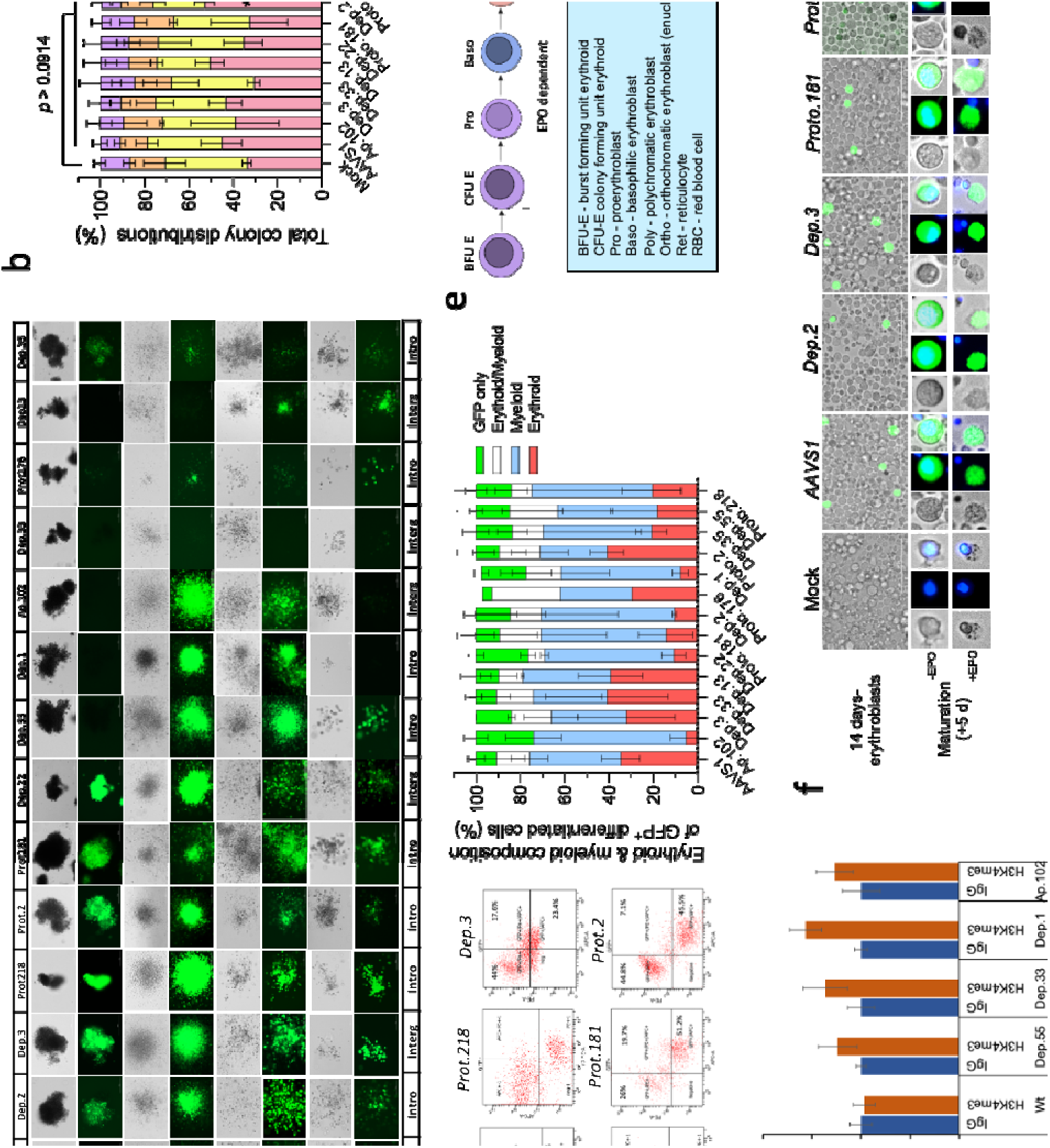
Multipotency of modified CD34+ HSPCs is retained after gene addition. **a** Colony forming units (CFUs) per targeted locus after 14 days in differentiation medium (bright field and epifluorescence images). The GSH locus designation is indicated on the top, intronic or intergenic category at the bottom, and CFU identity on the left (E, erythroid; G, granulocyte; GM, granulocyte–macrophage and M, macrophage). *AAVS1*, *Dep.2*, *Dep.3*, *Proto.218*, *Proto.2* and *Prot.181* GSHs allowed widespread transgene expression, whereas *Dep.55*, *Dep.1* and *Ap.102* restricted expression to CFU-G, -GM, and -M. Scale bar = 200 µm. **b** Total colony distributions per GSH-targeted cells. Mean ± s.d., *n* = 3 independent experiments. Two way ANOVA followed by Dunnett test * *P* < 0.05, ** *P* < 0.01. **c** Erythroid (CD235a^+^) and immune (CD69^+^) composition of GFP^+^ expressing cells. Representative FACS plots are shown (left). Mean ± s.d., *n* = 2-3 independent experiments. **d** CUT&RUN QPCR assay with primers amplifying the MND promoter of the transgene shows presence of H3K4me3 activation mark on the transgene inserted in *Dep.55*, *Dep.33*, *Dep.1* or *Ap.102* GSH sites of K562-derived megakaryocytes (*n* = 3 experimental replicates). **f** Erythroid maturation from *AAVS1*, *Dep.2*, *Dep.3*, *Prot.181*, *Prot.2*, and *Prot.218* unsorted cultures. *Upper*, 14 days-erythroblasts retain GFP+ signal. Scale bar 100 μm. *Bottom*, Enucleation occurs in presence (+EPO) but not in absence of human EPO (-EPO). Nuclear counterstaining with Hoechst 33342. Representative pictures of *n* = 2 independent human donors.

To confirm the multipotency of edited HSPCs, we analyzed the erythroid (CD235a^+^) and immune cell (with the general CD69^+^ cell surface marker) composition of the GFP^+^ differentiated population (Fig. 4c and 4d). The resulting compositions were similar among most analyzed groups relative to the *AAVS1* edited reference locus (Fig. 4d and 4e). Similar percentages of erythroid and immune cell phenotypes coexisted with small portions of unidentified GFP^+^ or GFP^+^/CD235a^+^/CD69^+^ triple labeled cells (Fig. 4e). Remarkably, the bias towards non-erythrocyte cell expression from *Dep.1*, *Dep.55*, and *Ap.102* GSHs was confirmed within the CD69^+^ fraction (Fig. 4e).

To characterize further the immune specific *eGFP* regulation, we established K562 cell-line models of *eGFP* integrated into *Dep.1*, *Dep.55*, *Dep.33,* or *Ap.102* GSHs. The K562 cell lines were then induced to erythroid (CD235a+) or megakaryocytic phenotype (CD61+) by treating with erythropoietin (hEPO) or phorbol myristate acetate (PMA), respectively (Supplementary Fig. 5a). K562-derived megakaryocytes may be considered as having an immune cell phenotype due to the recently described immune role of megakaryocytes^52^ and their separated origin from erythrocytes^53^. Thus, following induction with PMA, the megakaryocytic cells retained strong expression of the *eGFP* as expected (Supplementary Fig.5c). To determine whether epigenetic modifications were associated with transgene expression, CUT&RUN assays coupled with qPCR were performed. The results demonstrated that histone modifications showed enrichment of H3K4me3, but not H3K27me3, associated with the MND promoter of the transgene inserted in *Dep.1*, *Dep.55*, *Ap.102*, or *Dep.33* loci. Histones with H3K4me3 modifications are consistent with a sustained expression within immune cell lineages (Fig. 4d and Supplementary Figs. 5b, 5c and 6). In contrast to megakaryocytic induction, edited K562 cells treated with hEPO resulted in approximately 30% of induced cells positive for CD235a, an erythroid cell surface marker(Supplementary Fig.5d). Furthermore, the CD235a^+^ cell population was comprised of both GFP+ or GFP-cells. Similar results were obtained with erythroid induction with hemin (data not shown). DNA analysis confirmed the presence of properly integrated *eGFP* gene, thus GFP expression was most likely affected by epigenetic modifications; sequence analysis of the *eGFP* cassette would substantiate non-genetic basis for the GFP silencing (Supplementary Fig. 5d). These results are distinct from primary human HSCs in which the induced lineages were phenotypically homogeneous. Considering the leukemic origin of K562 cells, the model does not preclude the existence of immune specific GSHs observed after differentiation of modified CD34+ HSPCs.

### Persistent transgene expression in terminally differentiated erythroid cells

As the final step of erythroid differentiation comprises the erythroblast enucleation, we selected *Dep.2*, *Dep.3*, *Prot.218*, *Prot.2* and *Prot.181* groups for further erythrocyte maturation experiments with hEPO (Fig. 4g). Following the nucleofection of CD34^+^ HSPCs to target the mentioned GSH sites, the cells were expanded during 14 days in culture under erythroblast stimulating conditions (Fig. 2b). Next, the erythroblasts were exposed to hEPO for an additional five days of culture after which enucleation was confirmed in GFP^+^ cells, but not in non-hEPO treated cells (Fig. 4h). Of note, persistent transgene expression was observed in reticulocytes derived from all analyzed loci except for *Prot.218* (Fig. 4h) in which the transgene expression was silenced soon after exposure to hEPO (Fig. 4f).

Overall, the results support our approach for identifying useful GSH sites compatible with stable transgene expression, and in some cases tissue specific-expression helping to establish a concise catalog of sites for gene therapy of blood disorders.

## Discussion

This study demonstrated the feasibility of using evolutionary biology and comparative genomic approaches to identify prospective GSH in the human genome. Over the course of millions of years, parvovirus DNA genomes have integrated into the germline of multiple host species at genomic loci that tolerate DNA insertions. Of the previously 200 unique EPV insertions identified ^38^, we have mapped about half to orthologous sites in the human genome indicating that these loci are broadly conserved evolutionarily. Genomic loci were selected for CD34^+^ HSPC genome editing based on bioinformatic analyses which (i) determined precision of the EPV ortholog in the human genome (Fig. 1a, 1b, and Supplementary Table 1); (ii) predicted editability, and (iii) predicted transcriptional activity in human cells. In contrast to alternative approaches for identifying GSH candidate loci that rely on arbitrary assumptions considered to reduce risks of insertional mutagenesis ^16,19,21^, the methodology applied in the current study was intended to identify and characterize hematopoietic-compatible GSH loci. The criteria for establishing an ontogenic GSH (e.g., as in germline editing for transgenic mouse generation) are much different than the criteria used to establish a cell-, or lineage-, specific safe harbor, and as such, the “global” GSH is likely to be suboptimal for adult stem cells.

Following our approach, we evaluated and characterized novel genomic loci for gene therapy of blood disorders using primary human CD34^+^ HSPCs. However, further evaluation with *in vivo* models is necessary to demonstrate the ability of modified HSPCs for long-term engraftment of the bone marrow and reconstitution of blood cell compartments, ideally using a non-human primate model. As a first approximation, the *in vitro* results presented here indicate successful editing among the rare LT-HSC population (CD34^+^,CD90^+^) ^49^ even with relatively low numbers of CD34^+^ HSPCs used for editing. For these experiments, the number of nucleofected cells ranged from 2.5×10^5^ to 1.0×10^6^ for each GSH site being evaluated. In comparison, approved gene therapies against ADA-SCID (Strimvelis^TM^, Orchard Therapeutics, PLC), MLD (Libmeldy^TM^, Orchard Therapeutics, PLC) or β-thalassemia (ZYNTEGLO™, betibeglogene autotemcel, Bluebird Bio, Inc.) reportedly require up to 2.0-3.0×10^7^ autologous CD34^+^ HSPCs per kg of body weight, retroviral or lentivirally transduced, as the dose for infusion. However, indiscriminate lentivirus integration generates a heterogenous population of cells including cells that are killed outright, cells that lose stemness, and cells that do not contribute any therapeutic benefit. Therefore, logic supports the rationale that targeting an appropriate GSH with sufficient editing efficiency would yield a population of productive stem cells or CAR-T cells, and would reduce the number of apheresis rounds to obtain an appropriate number of cells for manufacturing.

Contrary to the large number of chimeric antigen receptor (CAR) T-cells reportedly required in cancer clinical trials (up to 1.0×10^8^ cells/kg) ^4,21,54^, expanding human CD34^+^ HSPCs by prolonged cell culture (>7 days) has been reported to detrimentally affect the long-term progenitor cells repopulating capacity ^47^. Recent studies of long-term HSPCs transplanted in patients with Wiskott-Aldrich syndrome or β-hemoglobinopathies estimated that as few as 98 edited HSPCs per 10^6^ infused CD34^+^ cells correspond to engrafting cells were sufficient to reconstitute gene-corrected granulocytes in circulation ^48^. This suggests that the more efficient the editing pipeline (which is partially dependent on efficient access to the genomic site), the greater the bioavailability of therapeutic cells in the infusion. If the novel GSHs are comparable to the highly efficient *AAVS1*, the up-scaling for clinical readiness (usually >5×10^6^ electroporated cells) might result in increased numbers of engraftable cells (i.e., modified LT-HSCs).

Although the GFP^+^ percentages of sorted cells were low for some targeted GSH loci (e.g. *Dep.1* or *Ap.102*), the values might easily be enhanced by cell cycle (S/G2) synchronization, screening additional gRNAs, or combined nucleofection/rAAV6 transduction as was demonstrated for *Ap.102*, *Dep.1*, *Dep.33* and *Dep.35* (Fig. 2h). In addition, persistent transgene expression in differentiated cells indicates that most of these GSH sites are compatible with long-term transgene expression making unnecessary the use of insulators to protect the transgene expression as discussed elsewhere ^21,55,56^. In this regard, a naturally occurring insulator in *AAVS1* ^57^ might explain the robust expression reported from this site (Fig. 2c and 4d) ^58^ suggesting that insulators artificially incorporated into the transgene cassette may improve the performance from targeted GSH sites. Whether the novel GSH sites contain insulators has not been determined, and the ssDNA templates used in this report were devoid of such elements. Nevertheless, constitutive reporter expression was maintained in immature as well as in both partially and terminally differentiated cells indicating that selected GSH loci are compatible with the long-term transgene expression in blood cells. Thus, these results establish a catalog of GSH with opportunities to engineer complex transgenes exploiting the permissiveness and tissue specificity of the novel GSH sites by including enhancers, insulators, IRES, T2A, or combinations that increase the transgene performance, or to engineer multigenic knock-ins that increase the traits of genetically modified hematopoietic cells. Remarkably, immune specific GSH sites introduce an additional level of safety restraining the transgene expression to the target cell population, particularly if intergenic GSHs are included as their modification conveys lower disturbances of the cellular transcriptome (Fig. 3a and 3b; Fig. 4). Similar tissue-specific GSHs are predicted to exist for all human tissue types.

In conclusion, this study established a concise catalog of 102 mapped GSH sites in the human genome with a sample of 17 loci yielding at least 9 GSH sites suitable for blood disorders therapy. Remarkably, the naturally occurring regulation from some GSH sites can be exploited for lineage restricted transgene expression. Thus, more GSH sites are likely to emerge from the remaining mapped loci with tissue specific-applications to modify other cell types.

## Methods

### Homology ascertainment relative to human genome

EPV positions were mapped to the orthologous human loci using Ensembl for comparative genomic alignments (GRCh38.p13) (Ensembl 2023). However, since the host species genome sequences accessible via Ensembl may be from different assembly versions than the reported EPV loci sequences the EPV coordinates within host genomes were acquired through BLASTN or BLAT analysis tool within the Ensembl suite. Then, using the Comparative Genomics options in Ensembl, the aligned host species and multiple mammalian genomes were analyzed to ascertain overall synteny and collinearity. Thus, the corresponding human genomic locations were determined. As some EPV host species were not available in Ensembl, BLASTN and EPV genomic flanking sequence data were used to locate the EPV integration site in closely related host species genomes that may or may not contain an EPV ortholog. Gaps in the host/human sequence alignment surrounding the EPV elements introduced another level of ambiguity in identifying sites in which the element may have been inserted.

### Selection of human genome loci to be evaluated as GSH

We focused on EPV loci that were mapped with a high degree of confidence on the human genome. As the experimental portion of this effort involved editing human primary CD34^+^ HSPCs, we evaluated the accessibility of chromatin for EPVs in this cell type. CD34^+^ HSPC data were retrieved from the Gene Expression Omnibus under accession GSE74912. ATACseq data were accessed with DolphinNext (Corces, et al.^39^) and chromatin accessibility was estimated through peak-calling with MACS2 with special attention for loci within 2 kb of an ATACseq peak.

### Cell culture and lipofection

Human embryonic kidney (HEK)-293T cells (ATCC CRL-3216) were cultured in 10% FBS DMEM (Gibco 11965-092), supplemented with penicillin (50 U per mL) and streptomycin (0.1 mg per mL) (Sigma P4458), at 37°C in 5% CO2 and saturating humidity. Transfection experiments on HEK-293T cells were performed in 6-well plates (Corning, NY). Briefly, one day prior to transfections, 3.5×10^5^ cells were added to each well. Cells were transfected with pX330 plasmid (2.5µg) using Lipofectamine 3000 (Invitrogen) following the manufacturer’s instructions. Forty-eight hours post-transfection genomic DNA (gDNA) was isolated with DNeasy Blood & Tissue Kit (Qiagen), and following concentration measurement, an aliquot of gDNA was used for PCR amplification of the targeted loci using locus specific primers to determine editing efficiency and specificity.

Human bone marrow-derived CD34^+^ stem and progenitor cells (HSPCs), obtained from healthy adult donors, were purchased from Stem Cell Technologies (SCT, Cambridge, MA 02142). Briefly, cryopreserved HSPCs were thawed and pre-stimulated for 48 h in StemSpanTM SFEM II medium (SCT 09655) with Expansion supplement (SCT 02691) and 1 μM UM729 (SCT 72332), and incubated at 37° C in 5% CO_2_ and saturating humidity atmosphere. Following electroporation, the CD34^+^ HSPCs were returned to the same incubation conditions for additional 72 h before further downstream use.

Erythroleukemia K562 (ATCC CCL-243) cells were maintained in 10% Iscove’s modified Dulbecco’s medium(IMDM) (10-016-CV, Corning Life Sciences) supplemented with penicillin (50 U per mL) and streptomycin (0.1 mg per mL) (Sigma P4458), at 37° C, 5% CO2 and saturating humidity. Following electroporation, the cells were returned to the growth medium and incubated for 5 days before cell sorting. Sorted cells expressing GFP were expanded under the same incubation conditions for use in subsequent experiments.

### Guide RNAs for CRISPR/Cas9 gene editing

Single guide RNAs (gRNAs) were simultaneously engineered with CHOPCHOP, CRISPOR and CRISTA designing tools. Three to five highly scored gRNAs with minimal off-target sites that were common to the three gRNA design programs, were chosen per candidate GSH locus. Synthetic oligonucleotides incorporating the *gRNA* sequences were ligated into pX330 plasmid (Addgene, Watertown, MA 02472) to co-express *Cas9* and *gRNA*. Indel formation of the target site was used as an indicator for editing efficiency in HEK-293 cells. Forty-eight hours post-transfection into HEK293T cells, genomic DNA was isolated (Qiagen), and PCR amplified into 600 bp products (Takara). Indel frequencies were determined through Sanger data-decomposition using the TIDE analysis tool (tide.nki.nl/). Indel efficiencies were compared to that provided for a widely reported *AAVS1* gRNA (Supplementary Table 3). Thus, the most efficient gRNA per locus was selected for downstream experiments in CD34^+^ HSPCs. Next, a chemically synthesized version of each gRNA, modified at both termini with 2’-O-methyl-3’-phosphorothioate, was purchased as a single guide molecule from IDT (Integrated DNA Technologies, IA) and complexed with ArchiTect™ Cas9 nuclease (Stem Cell Technologies) for editing experiments in primary human CD34^+^ HSPCs.

### Single-stranded DNA template preparations

Single-stranded DNA donor template (ssDNA) was synthesized from a plasmid carrying 300 bp homology arms matching the corresponding GSH site flanking the *eGFP* expression cassette. Briefly, an expression cassette containing the *eGFP* open reading frame (ORF) regulated by the MND promoter and rabbit β-globin poly(A) signal was constructed by using conventional molecular biological techniques. Homology arms were obtained by PCR from HEK-293T genomic DNA and a plasmid template was generated per candidate GSH site by Gibson assembly (New England Biolabs) using fragments synthesized by Q5 High Fidelity Taq polymerase (New England Biolabs). Finally, 5’-phosphorylated PCR primers were used to amplify the 2.0 kb eGFP expression cassettes flanked with 300 bp homology arms (Supplementary Table 6). To produce single-stranded template DNA, the phosphorylated reverse strand from each substrate was digested by sequential strandase-treatment to yield ssDNA (Takara 632666) which was column-purified and resuspended in nucleases-free water (Gibco). Plasmid templates were eliminated using DpnI treatment (NEB R0176) prior to the ssDNA synthesis..

### Electroporation

Prior to nucleofection of CD34^+^ HSPCs, ArchiTect™ Cas9 nuclease (SCT) and the corresponding gRNA were combined (1:2.5 molar ratio) to form a ribonucleoprotein complex at 25° C for 20 min and placed at 4° C until used for nucleofection reactions. In parallel, CD34^+^ HSPCs pre-stimulated for two days were washed in 50 mL PBS at 37° C and resuspended in P3 nucleofector solution at RT (Lonza, V4XP-3032). Approximately 2.5×10^5^ CD34^+^ HSPCs were mixed with gRNA:Cas9 complexes and 3 µg ssDNA donor template (20 μL) in nucleofector strip format (Lonza). Electroporation was performed in Unit X, Lonza 4D nucleofector, using the DZ-100 program. After nucleofection, cells were incubated for 10 min at room temperature before adding the StemSpan™ SFEM II medium with Expansion supplement and 1 μM UM729 for a 2 day-recovery time. After the recovery period, the cells were sorted by flow cytometry and GFP^+^ cells were collected. Similarly, erythroleukemia K562 cells were washed in 50 mL PBS at RT and approximately 1×10^6^ cells were resuspended in SF solution (Lonza V4XC-2032) and electroporated in presence of pre-assembled Cas9:gRNA complexes plus 12 µg ssDNA (100 µL format) in a Lonza 4D-nucleofector using the FF-120 program.

### AAV6 transduction

Recombinant adeno-associated virus type 6 (rAAV6) vectors were obtained from the University of Massachusetts Chan Medical School Viral Vector Core facility. Cryopreserved CD34^+^ HSPCs were thawed, transferred to tissue culture plates, and incubated in StemSpan^TM^ SFEM II medium for a 2 days pre-stimulation period under conditions described above. Following pre-stimulation treatment, 2.5×10^5^ CD34^+^ cells were electroporated in the presence of Cas9:gRNA complexes, followed by a recovery period of 15 min at RT. Next, cells were collected and transduced with rAAV6 at a multiplicity of infection (MOI) of 200,000-300,000 gc/cell (genome copies) in a 15 mL conical tube (Corning). Cells were transferred to a 6-well culture plate (Corning) at a density of 1×10^5^ cells/mL and incubated for 2 days to recover from the treatment before downstream analysis.

### Colony Forming Unit assay (CFU)

Between 300 and 1500 sorted GFP^+^ cells were seeded in 1 mL methylcellulose hematopoietic progenitor cell differentiation medium (MethoCult, SCT H4435) and transferred to 6-well SmartDish plates (SCT 27371). Differentiation was induced during 14 day incubation at 37°C, 5% CO_2_ and saturating humidity before scoring the Colony Forming Units (CFUs) by brightfield microscopic inspection (Axiovert 135 microscope, Zeiss) and fluorescent images were obtained with fluorescent microscopy (LionHeart FX,BioTek). After colony scoring, the entire plate of cells were recovered from methylcellulose by diluting in PBS, washed and resuspended in ice cold 1% FBS in PBS before the staining for flow cytometry analysis as described above.

### Flow cytometry

After recovering for 2 days, nucleofected CD34^+^ HSPCs were washed once in PBS and resuspended in ice cold phenol red-free-IMDM (Gibco) supplemented with 1% FBS (Gibco) and processed using fluorescence activated cell sorting (FACS). Non-transfected HSPC samples were used for gating GFP^+^ signals. Propidium iodide (Sigma) was included for viability determinations then, GFP^+^ cells were collected and cultured for 3 additional days in StemSpan^TM^ SFEM II, CD34^+^ Expansion supplement and 1 μM UM729. The fold-expansion was determined by dividing the number of cells after three days of culture by the number of GFP^+^ cells seeded immediately after sorting. Thus, the total timeline for nucleofection experiments starting with cryopreserved, immature CD34^+^ HSPCs spanned 7 days. For the analysis of immature populations (LT-HSC and MPPs), cells were cultured for five days post-nucleofection, washed in PBS and then resuspended in ice cold PBS supplemented with 1% FBS. Subsequently, cells were incubated in presence of TruStain FcX^TM^ (BioLegend) before staining with antibodies against CD34 (BioLegend 343625), CD90 (BioLegend 328114) and CD38 (BioLegend 303506). Samples were fixed in ice cold 2% paraformaldehyde (PFA) for 10 min, washed and resuspended in phenol red free-IMDM. The GFP^+^/CD34^+^ population was gated to explore the modified immature cell populations. For cell lineage determinations (erythroid and immune), cells were recovered by diluting the methylcellulose in PBS and resuspending the cells in ice cold 1% FBS in PBS. The Fc receptors were blocked with Human TruStain FcX^TM^ (BioLegend) before staining with antibodies against CD235a and CD69 for 20 min at 4°C in the dark. All antibodies used in this work can be consulted in Supplementary Table 4. Following the incubation, the cells were washed in ice cold PBS and resuspended in ice cold 2% PFA for 10 min, washed once again with ice cold PBS, and finally resuspended in 1% FBS phenol red-free IMDM (Gibco) before the analysis. For experiments with differentiated K562 cells, the cells were stained with antibodies against CD235a (erythroblastic) or CD61 (megakaryocytes) surface markers as described above except that fixation with 2% PFA was omitted as the samples were processed immediately for CUT&RUN epigenetic modification analysis.

### Erythroid differentiation

Nucleofected CD34^+^ HSPCs were maintained in SFEM II cell culture medium with Erythroid supplement (SCT) to induce erythroblast expansion during 14 day incubation. Medium changes were performed at days 7 and 11 of culture. At day 14, cells were transferred into SFEM II supplemented with 3 U/mL hEPO (SCT) and 3% AB human serum (Sigma) for an additional 5 days of culture. Cell nuclei were stained with Hoechst 33342 (ThermoFisher) prior to microscopy imaging.

### Erythroblastic and Megakaryocyte induction of K562 cells

K562 cells were induced to erythroblastic cells by exposure to 3U/mL hEPO (SCT 78007) or 50 ìM hemin (Sigma). Megakaryocytic cells were obtained with 80 nM phorbol 12-myristate 13-acetate (Invivogene). Cellular phenotypes were identified by cell surface markers using anti-CD235a antibody (BioLegend 349105) for erythroblastic and anti-CD61 antibody (BioLegend 336406) for megakaryocytic cells and flow cytometry. The sorted cells were immediately processed for CUT&RUN experiments.

### Droplet digital PCR

The integration efficiency was determined by ddPCR of genomic DNA extracted from sorted GFP^+^ cells of 14 days post-nucleofection. Briefly, genomic DNA isolated from the cells (Qiagen) was amplified by ddPCR in the presence of a primer/FAM probe set specific for the integrated transgene and a primer/HEX probe set specific for a reference region within the same unaltered locus (Supplementary Tables 7, 8 and 9). Reaction mixtures consisted of 1x ddPCR Supermix for probes (No dUTP) (Bio-Rad), primer/probe mixtures (900 nM primer/250 nM probe) and up to 60 ng gDNA per reaction. The thermocycler was programmed as follows: initial denaturation (95°C for 10 min with ramp rate of 2°C/sec), followed by 60 cycles of denaturation (94°C for 30 sec, ramp rate of 2°C/sec), annealing (55°C for 1 min ramp rank of 1°C/sec), and extension (72°C for 2 min, ramp rate of 1°C/sec), and a terminal denaturation step (98°C for 10 min), followed by cooling (4°C). Droplet Reader and Quantasoft analysis software (Bio-Rad) were used to analyze the fluorescence using 1D and 2D plots for thresholding FAM and HEX positive droplets. Ratios of FAM to HEX were used to calculate the abundance of edited alleles.

### RNA-seq

Sorted GFP^+^ HSPCs were cultured for three days in StemSpanTM SFEM II medium (SCT) before being washed in PBS. Subsequently, the cells were mixed with five volumes of RNA*later* solution (Thermo Fisher AM7020) and stored at -80° C until processing for RNA-seq analysis. For library preparation, the mRNA fraction was prepared by poly(A)+ selection (Illumina). Libraries were sequenced as paired end 150 bp reads on an Illumina HiSeq 3000 with a sequencing depth of approximately 20-30 million per sample. Quality control metrics were confirmed in FASTQ files through FastQC (<u>bioinformatics.babraham.ac.uk/projects/fastqc/</u>). Adapter sequences were removed from the reads and mapped to the *Homo sapiens* reference genome (GRCh38) available on ENSEMBL using STAR aligner v.2.5.2b. Differential expression analysis was performed by analyzing un-normalized read counts using DESeq2 and applying “donor” as a cofactor. Transcripts with an adjusted p-value < 0.05 and absolute log_2_ fold change > 1 were designated differentially expressed genes (DEGs) as d. Heatmaps with log_2_ normalized read counts were generated with R Studio, clustering genes according to their regulation changes. Gene ontology (GO) analysis was performed with the iPathwayGuide platform of Advaita Bioinformatics (<u>advaitabio.com</u>).

### CUT&RUN Assay

Chromatin samples were processed using the EpiCypher CUTANA™ ChIC/CUT&RUN kit (EpiCypher 14-1048), according to the manufacturer’s protocol. Briefly, approximately 500,000 cells per sample were collected by centrifugation, resuspended in PBS, then re-centrifuged and then coupled to activated Concanavalin A beads. The cells were incubated with antibodies overnight at 4°C. Approximately 0.5ug of an antibody was used per sample, including IgG negative control antibody (EpiCypher 13-0042), H3K4me3 (EpiCypher 13-0041), and H3K27me3 (Active motif 39055). For permeabilization, 0.01% of digitonin was used in the buffers. Unbound antibody was removed by washing 2x, the cells were incubated with protein A and protein G - micrococcal nuclease fusion (pAG-MNase) to recover the histone - DNA complex. MNase was activated by adding 1mM CaCl2 and incubated at 4°C for 2 hours, releasing the chromatin immune complexes into the supernatant. The MNase reaction was terminated using a stop buffer master mix and to remove RNA from the sample. The released DNA fragments were recovered and size selected using SPRIselect beads and eluted in 15ul of elution buffer for subsequent qPCR analysis. Genomic DNA extraction was extracted from the input sample (1% of total cells) and the enrichment of immunoprecipitated DNA was calculated using percent input method formula - 100*2^(Adjusted Input – CT (IP). The final values were plotted as fold enrichments over IgG negative control.

### Statistical analysis

All the experiments with primary human cells were independently performed at least three times for statistical analysis. Cells were harvested from different anonymous, healthy BM donors each time. Data represents mean ± SD. Statistical analyses were performed by ANOVA with Dunnett test post hoc. Significant and non-significant *P* values are indicated on the Figures and the main text. Results were considered significant when *P* < 0.05. When less than 3 experiments were performed for descriptive results, an opportune mention is indicated on the figure legends.

## Supplemental Information

Supplemental Documents SX – Locations of EPVs on Human Genome (eve-meta-hg38)

## Data Availability

Sequence data generated for this study have been deposited in the National Center for Biotechnology Information Sequence Read Archive under BioProject PRJNA1022496.

## Acknowledgements

This work was supported in part by funding from the Association Monégasque Contre les Myopathies (RMK), and the Bill & Melinda Gates Foundation (OPP1202116 to RMK).

## Author contributions

R.M.K and M.Q-R designed experiments and analyzed data. M.Q-R. performed the experiments. S.L., M.A.C., and R.G analyzed genomes to map EPVs to the human genome. M.Q-R and M.A.C analyzed RNA-seq data. K.M.P. performed the CUT&RUN experiments. R.M.K. conceptualized and obtained funding. M.Q-R drafted the manuscript. All the authors revised and approved the final version.

---

**Supplementary Table 2.**
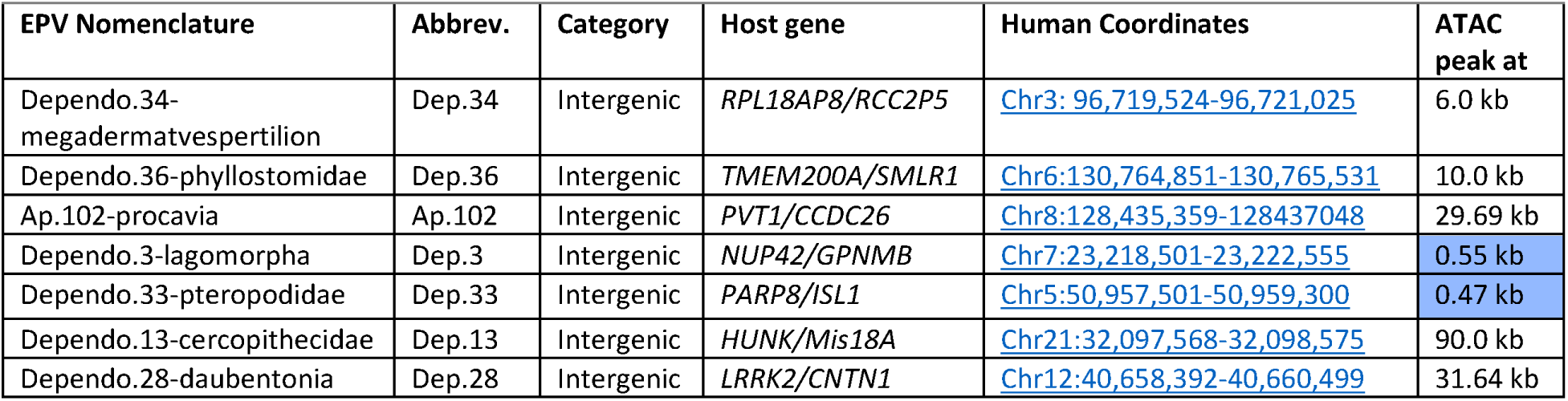
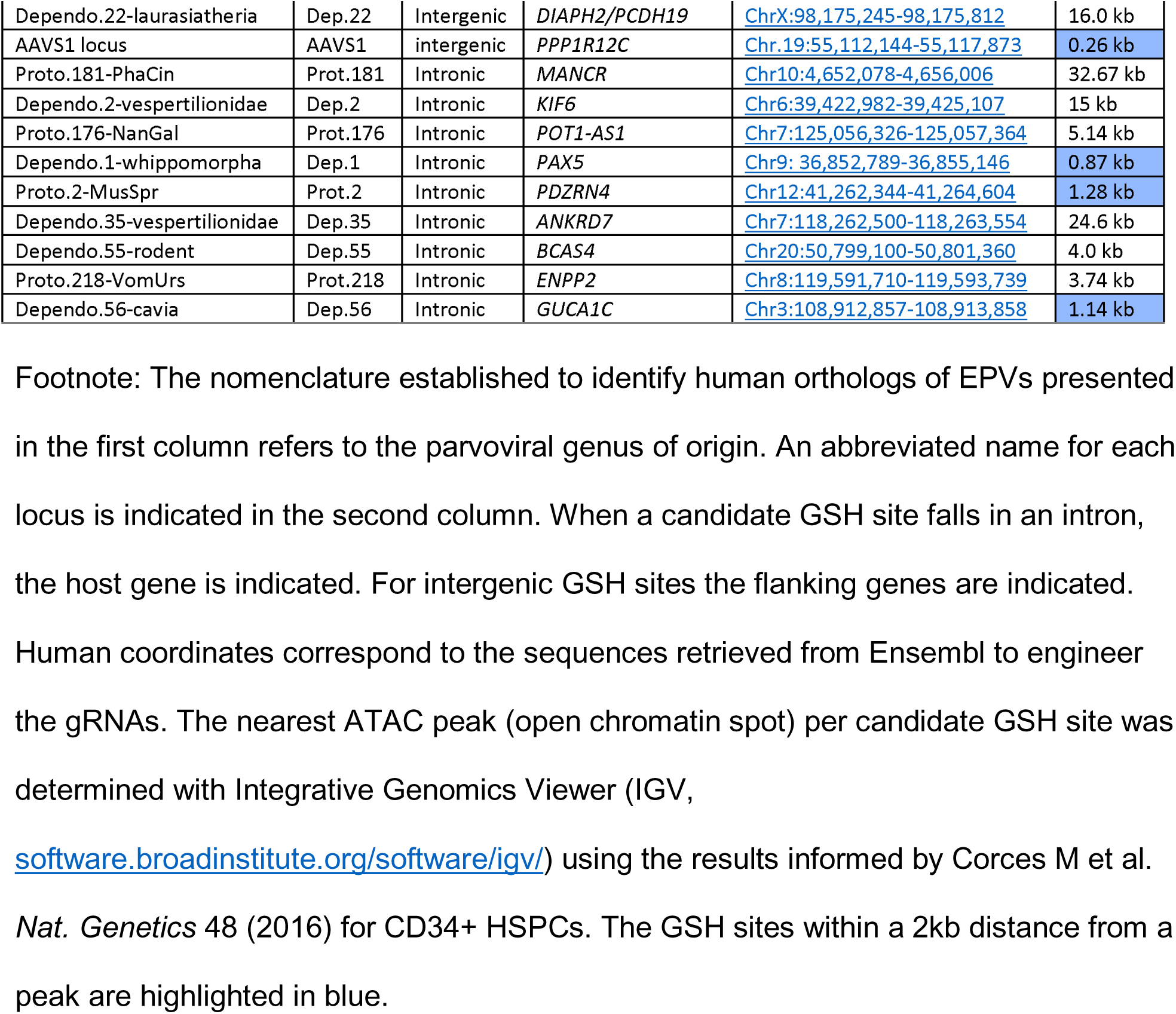
Human GSH sites experimentally assessed in primary CD34+ HSPCs.

**Supplementary Table 3.**
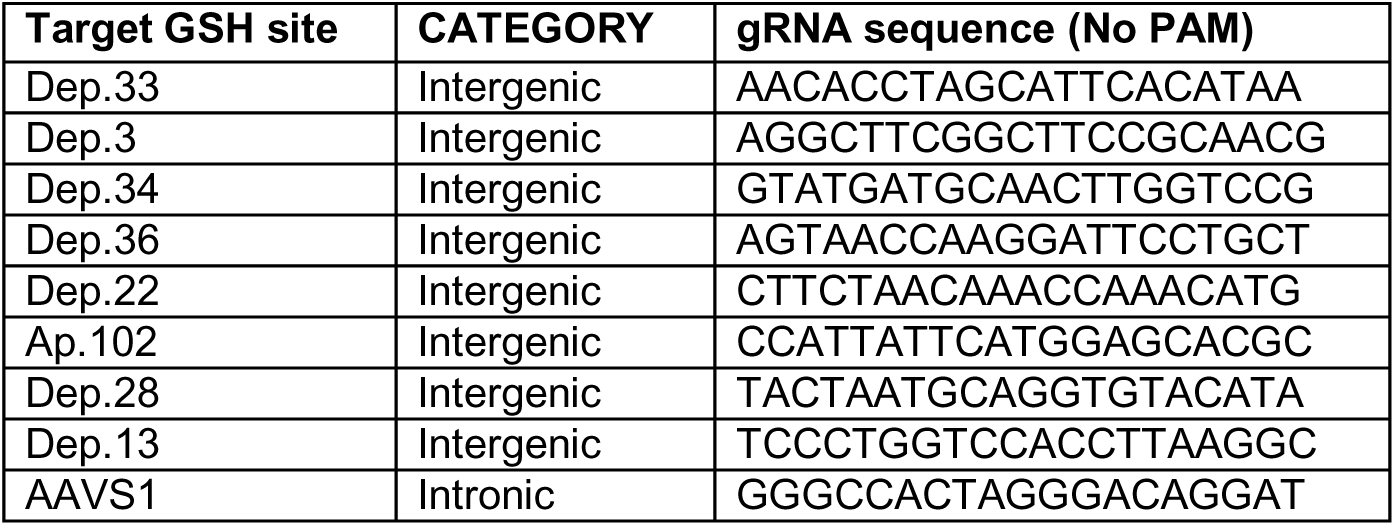
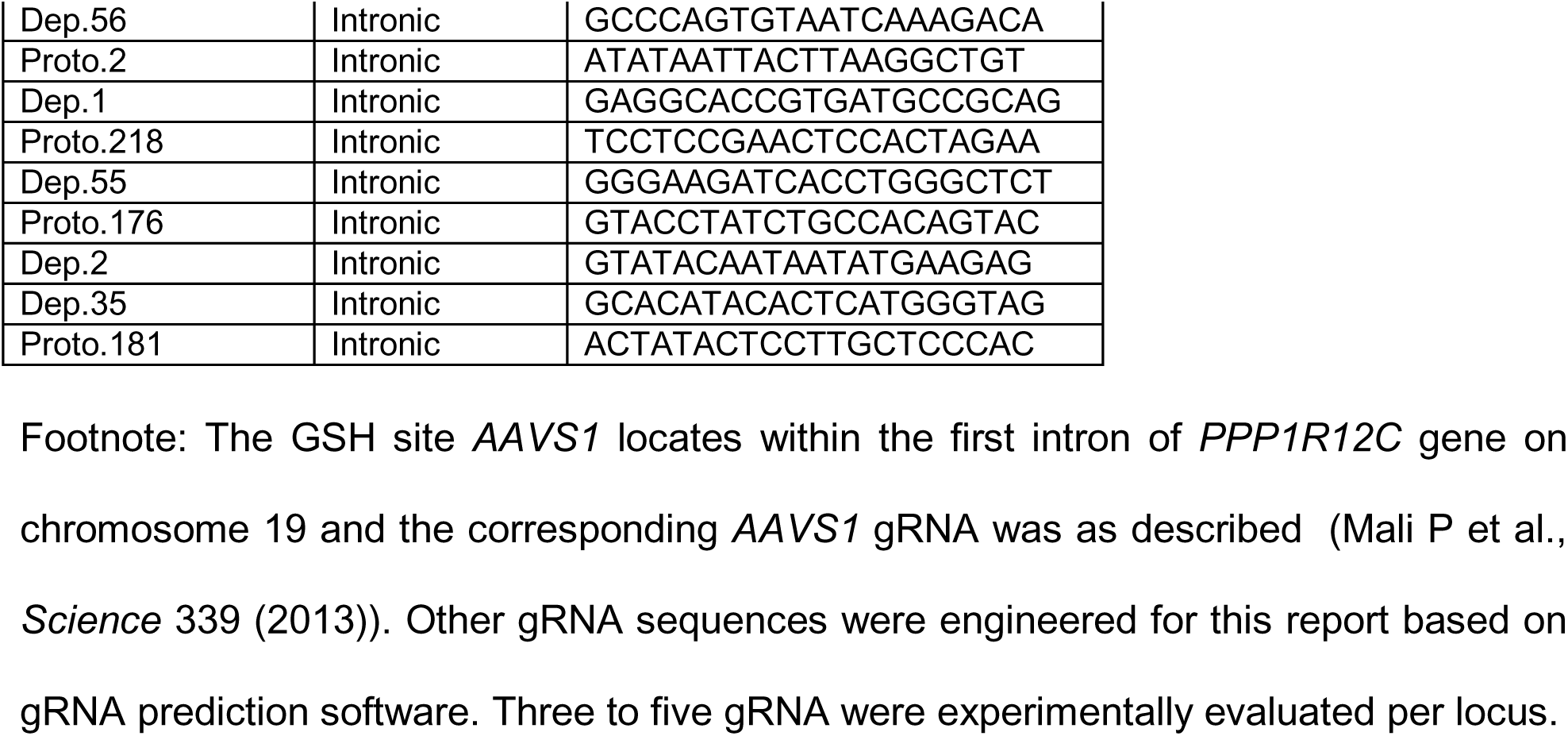
Guide RNAs selected per candidate GSH site.

**Supplementary Table 4.** Candidate GSHs showing persistent transgene expression during differentiation experiments of blood cells.

| GSH | Category | GFP<sup>+</sup> % | P value | ATAC peak at |
| --- | --- | --- | --- | --- |
| *Dep.3* | Intergenic | 2.07 ± 0.54 | 0.8495 | 0.55 kb |
| *Dep.22* | Intergenic | 1.32 ± 1.14 | 0.0258 | 16.0 kb |
| *Ap.102* | Intergenic | 0.62 ± 0.41 | < 0.0001 | 29.69 kb |
| *Dep.2* | Intronic | 3.23 ± 0.86 | 0.8938 | 15.0 kb |
| *Prot.181* | Intronic | 1.35% ± 1.05 | 0.0473 | 32.67 kb |
| *Dep.1* | Intronic | 0.54 ± 0.36 | < 0.0001 | 0.87 kb |
| *Prot.2* | Intronic | 1.0% ± 0.51 | 0.0027 | 1.28 kb |
| *Dep.55* | Intronic | 1.56 ± 0.53 | 0.1076 | 4.0 kb |
| *Prot.218* | Intronic | 1.62 ± 0.53 | 0.1961 | 3.74 kb |

**Supplementary Table 5.** Reagents used during this research.

| Reagent | Catalog number |
| --- | --- |
| Anti-human CD34 | BioLegend #343625 |
| Anti-human CD69 | BioLegend #310909 |
| Anti-human CD235a | BioLegend #349105 |
| Anti-human CD90 (Thy1) | BioLegend #328114 |
| Anti-human CD38 | BioLegend #303506 |
| TruStain FcXTM | BioLegend #422302 |
| IgG control antibody | EpiCypher #13-0042 |
| Anti-H3K4me3 | EpiCypher #13-0041 |
| Anti H3k27me3 | Activemotif #39055 |
| SFEM II medium | SCT #09655 |
| CD34+ Expansion Supp | SCT #02691 |
| UM729 | SCT #72332 |
| Methocult | SCT# H4435 |
| Erythroid Expansion Supp | SCT # 02692 |
| Huma EPO | SCT # 78007 |
| P3 primary solution | Lonza V4XP-3032 |
| ArchiTect™ Cas9 nuclease | SCT #76004 |
| ddPCR Supermix for probes (No dUTP) | BioRad #1863024 |
| ssDNA Production System | Takara #632666 |

**Supplementary Table 6.**
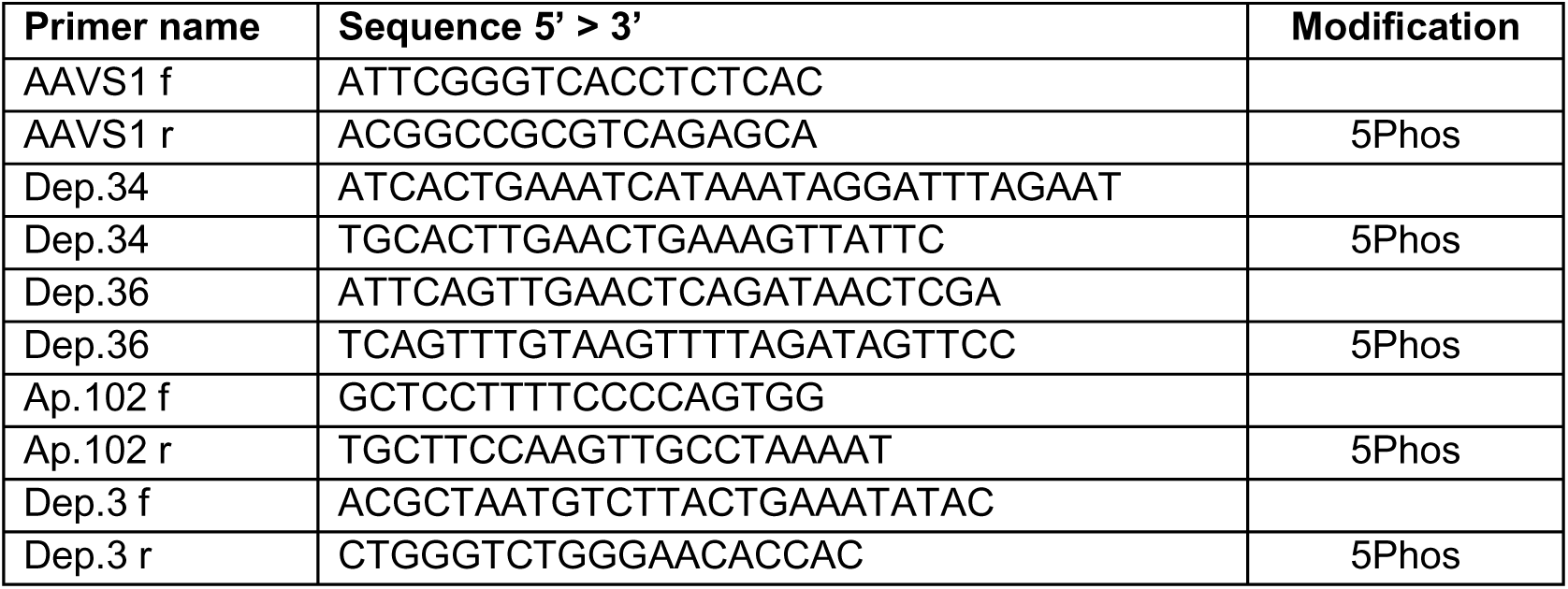
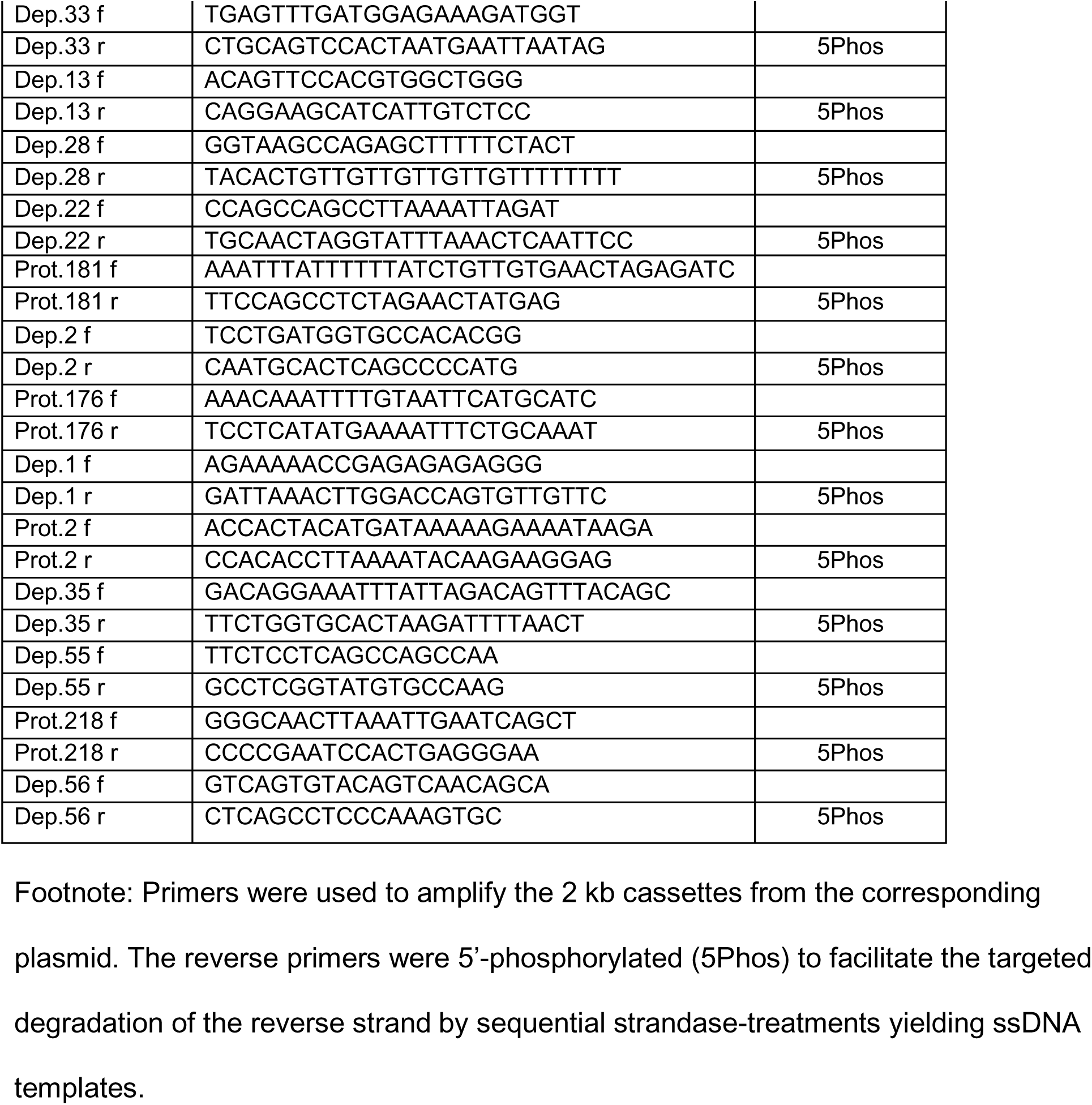
Primers used for ssDNA synthesis.

**Supplementary Table 7.**
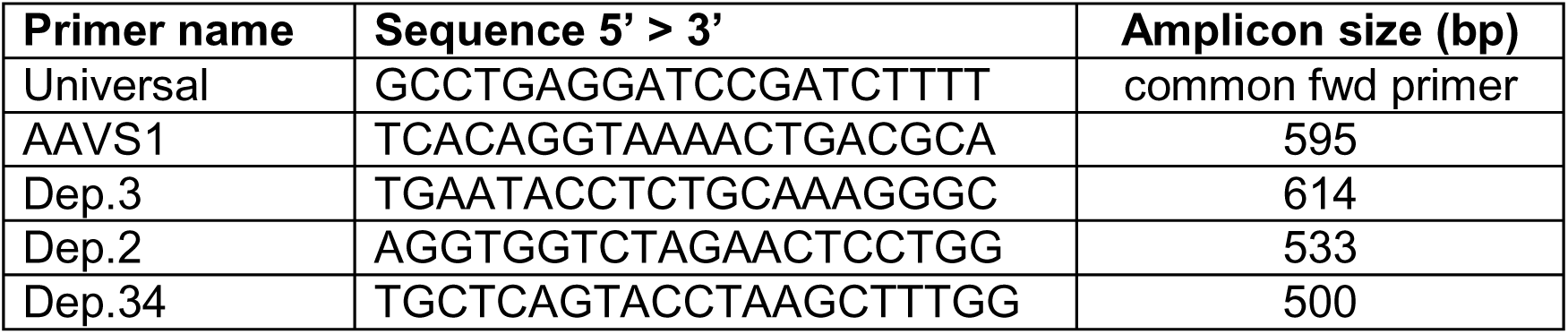
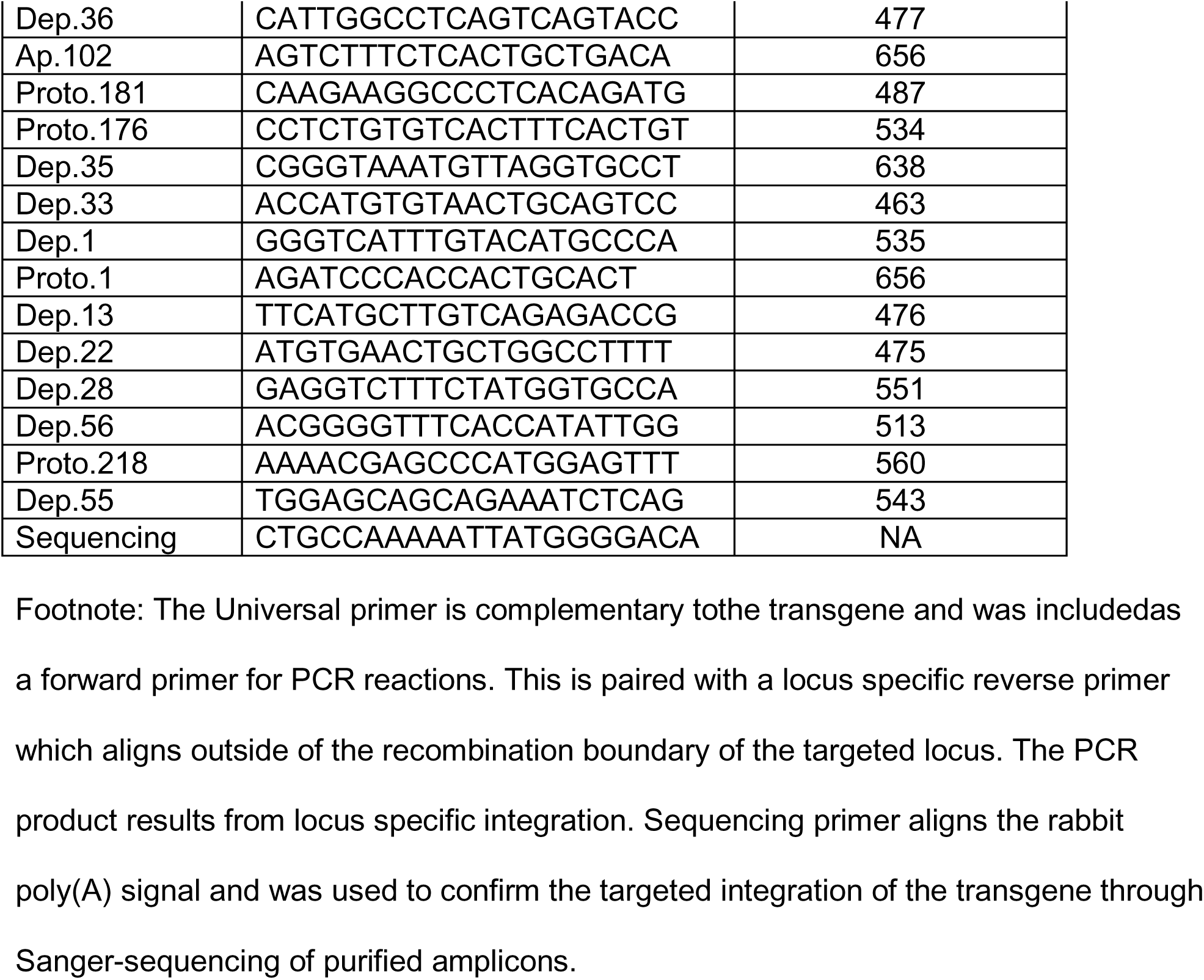
Primers to determine GSH-specific integration through ddPCR.

**Supplementary Table 8.**
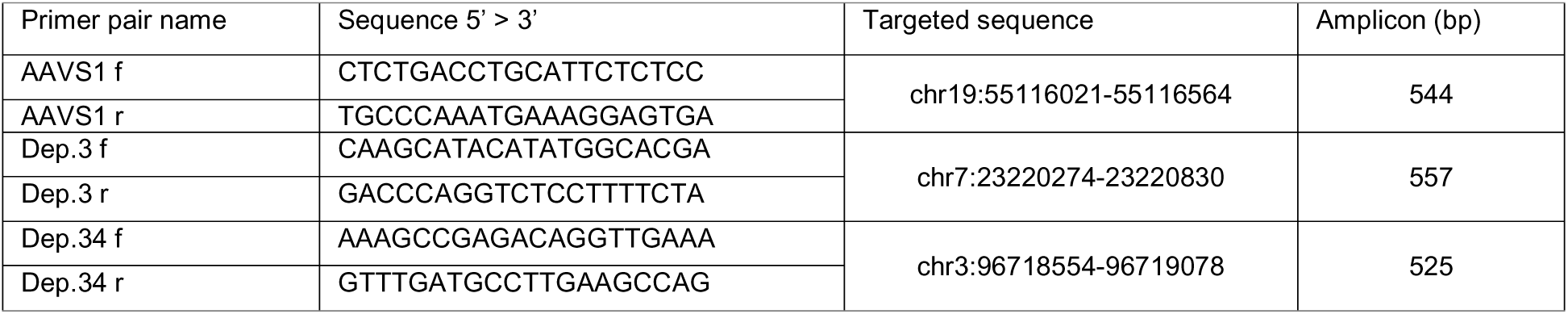
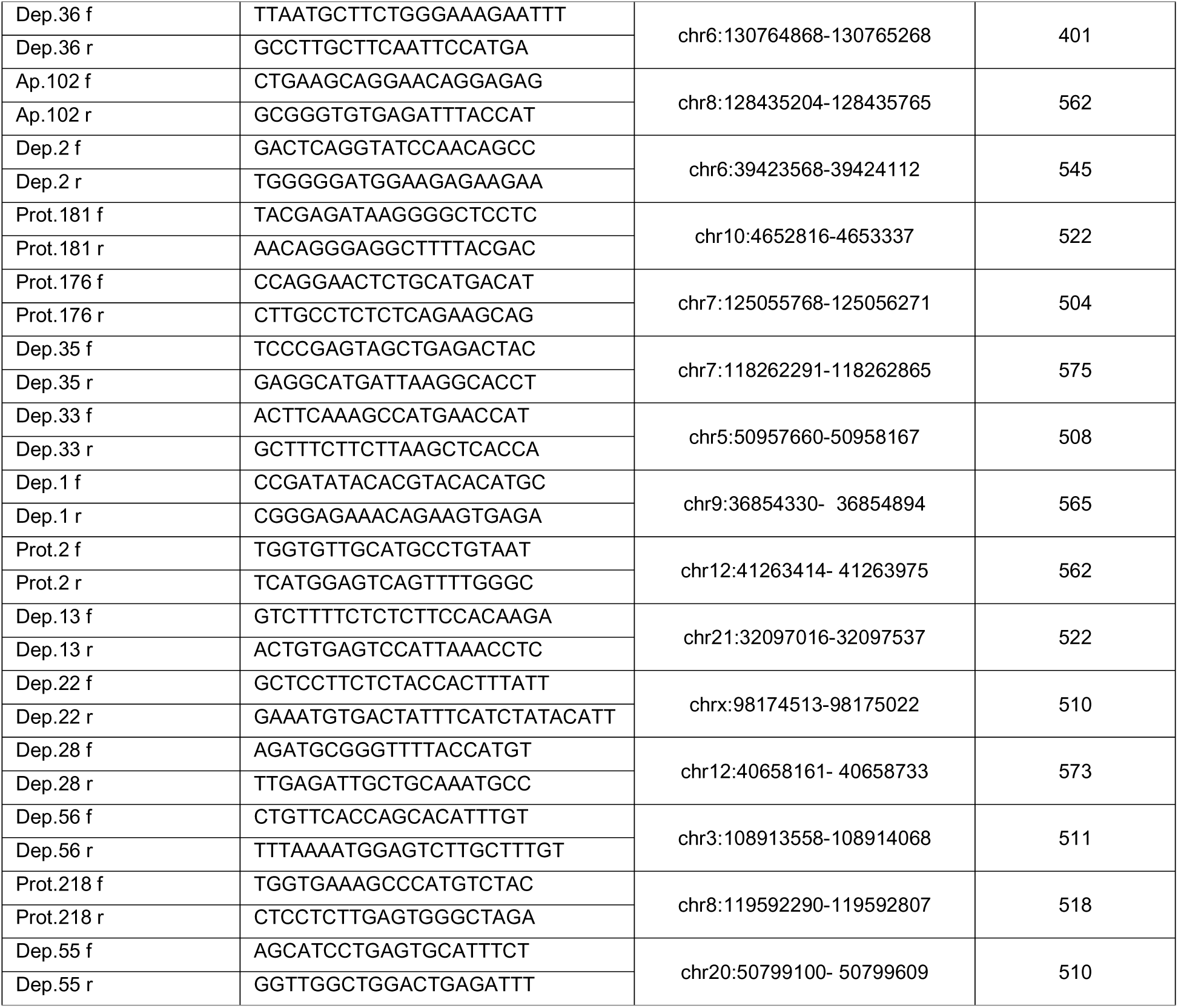
Reference primers recognizing an unedited region of the candidate GSHs.

**Supplementary Table 9.**
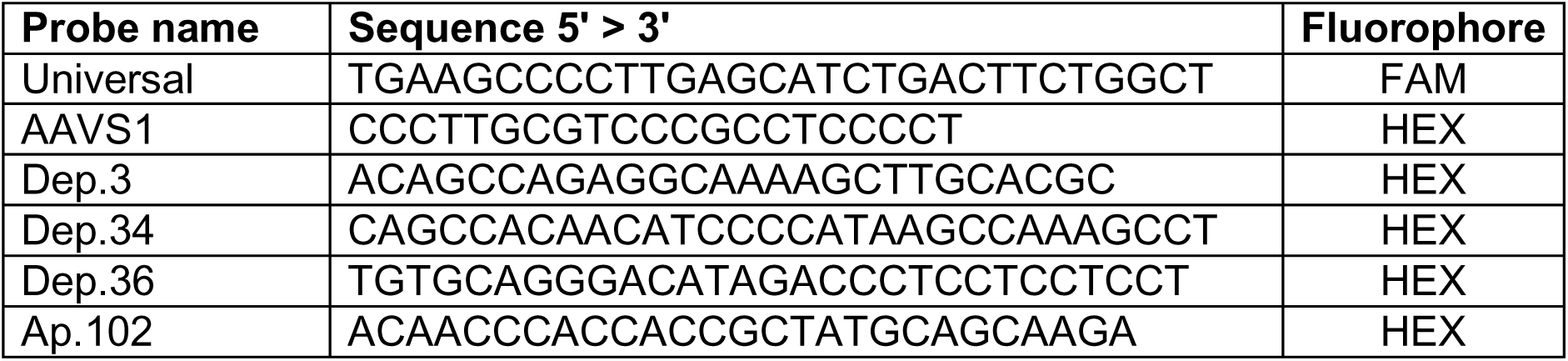
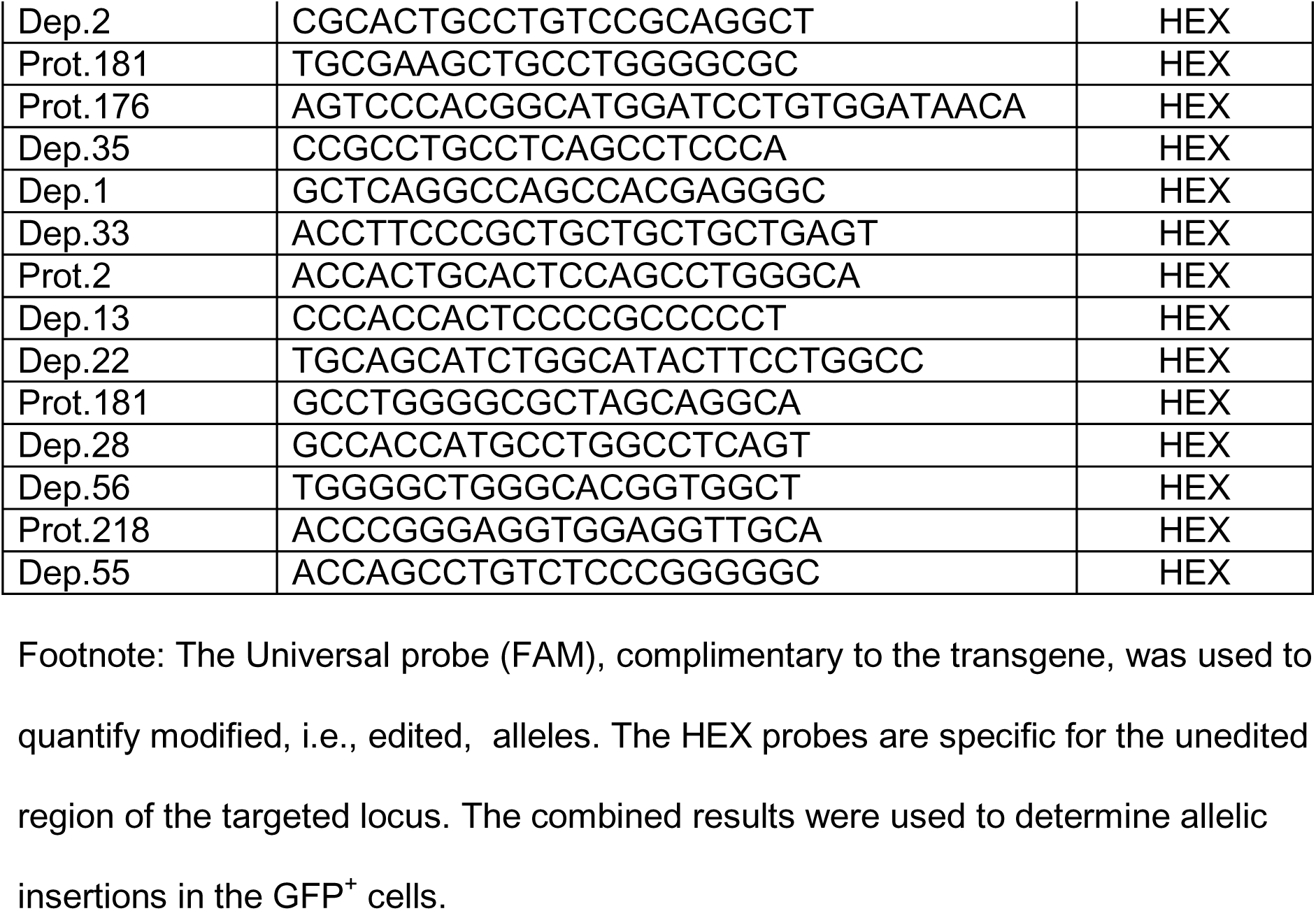
Fluorescent probes for ddPCR.

**Supplementary Table 10.**
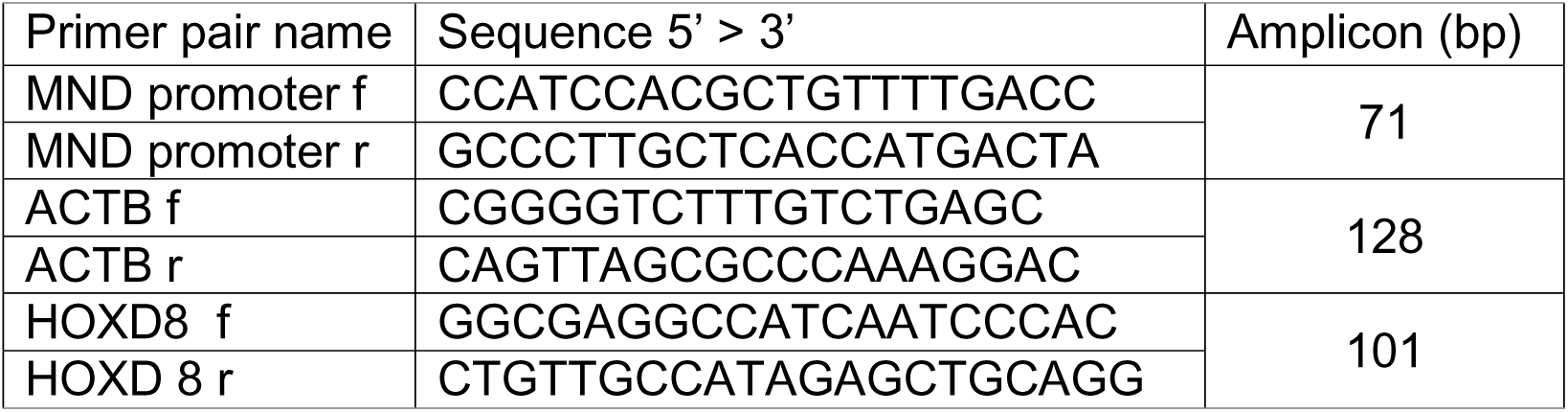
Primers for CUT&RUN qPCR.

**Fig. S1.**
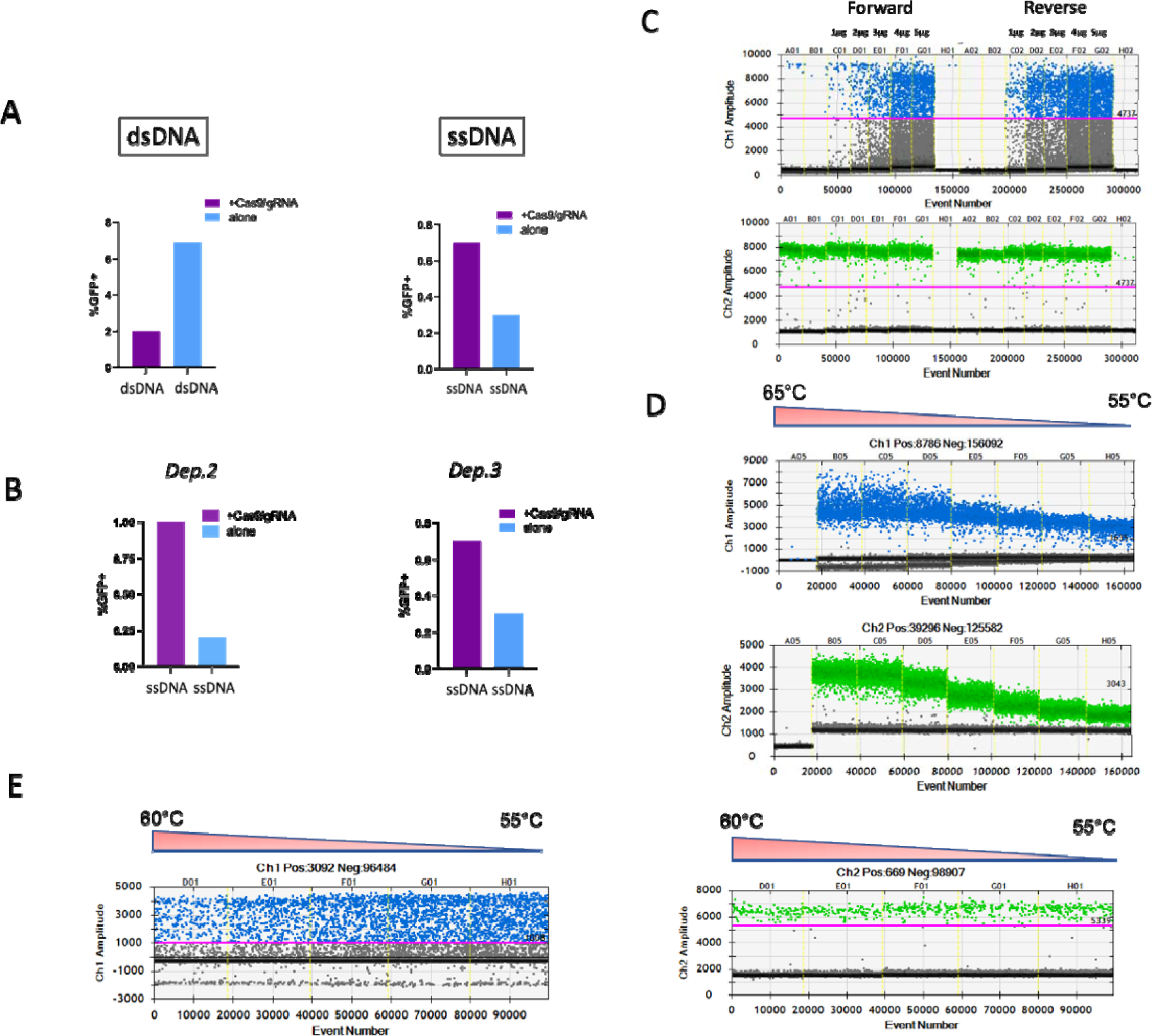
Comparison of dsDNA and ssDNA donor templates for genome editing of human CD34+ HSPCs. A) Two days pre-stimulated CD34^+^ HSPCs were transfected with dsDNA (HITI design) or ssDNA donor templates (HR), both carrying the same *eGFP* cassette, in presence or absence of CRISPR/Cas9 complexes specific for *Dep.2* locus. Two days after the nucleofection the GFP^+^ cells were sorted by FACS. Nucleofection with dsDNA alone (i.e., in absence of CRISPR/Cas9 complexes) resulted in a greater number of GFP^+^ cells than cells including CRISPR/Cas9 complexes. In contrast, cells treated with ssDNA alone showed less percentage of GFP^+^ cells in absence of CRISPR/Cas9 complexes indicating that ssDNA itself prevents GFP expression from non-integrated templates and suggests that the likelihood of random integration of ssDNA occurs infrequently, perhaps due to reduced stability of the ssDNA template. B) The same phenomenon was observed independently of the targeted locus (*Dep.2* and *Dep.3* are shown). C) Nucleofection of dsDNA templates also affect the quantification of edited cells through ddPCR as shown by 1D plots. The GFP^+^ cell populations, processed by ddPCR, displayed the same rain pattern with increasing amounts of dsDNA (1-5 μg). The primer sets to detect the transgene inserted in forward or reverse orientation (following the HITI approach) produce a similar rain effect (blue droplets) in Quantasoft. Conversely, primers recognizing an unedited region (green droplets) produce defined droplet clusters. D) 1D plots from CD34^+^ HSPCs transfected with ssDNA templates targeting *Dep.2* locus. Primers amplifying the junction regions (as described in main text) and a reference region generate defined clusters for quantitation even at different temperatures. E) Delivery of templates through rAAV6 produces as well an increased noise/background (rain effect) as shown by targeting of the *Dep.13* locus. Notice that the reference amplimer (right) did not show changes under the same temperature gradient.

**Fig. S2.**
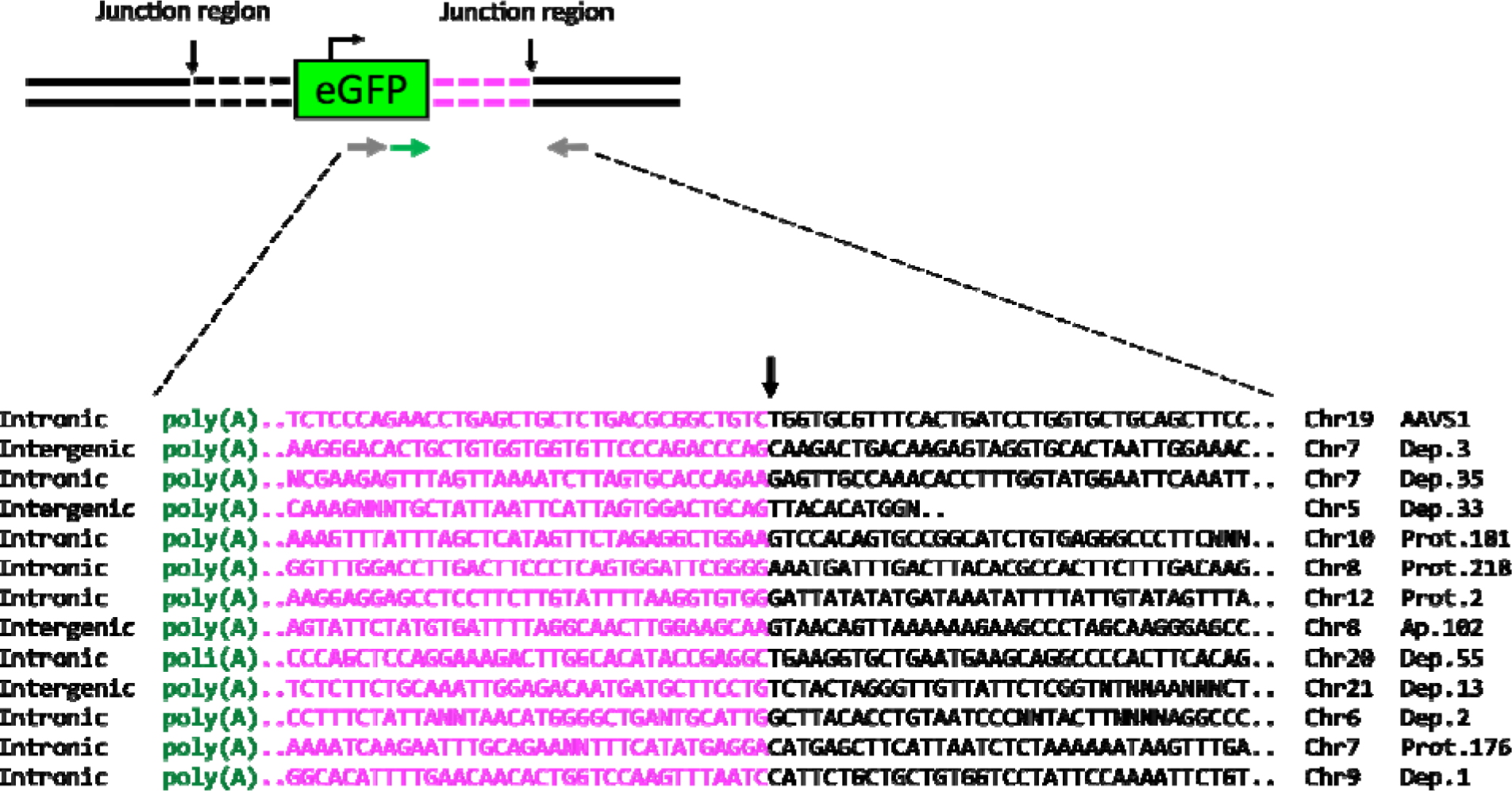
GSH-specific integration using ssDNA templates. Amplicons obtained from edited primary HSPCs (oligos available in Supplementary Table 6) were gel purified and Sanger-sequenced with a common primer annealing the rabbit poly(A) signal. All amplimers started with the poly(A) sequence included in the transgene (green) followed by the right homology arm used for targeted recombination (magenta). The arrow indicates the junction between the right homology arm and the corresponding targeted locus (black) showing continuity among the sequences.

**Fig. S3.**
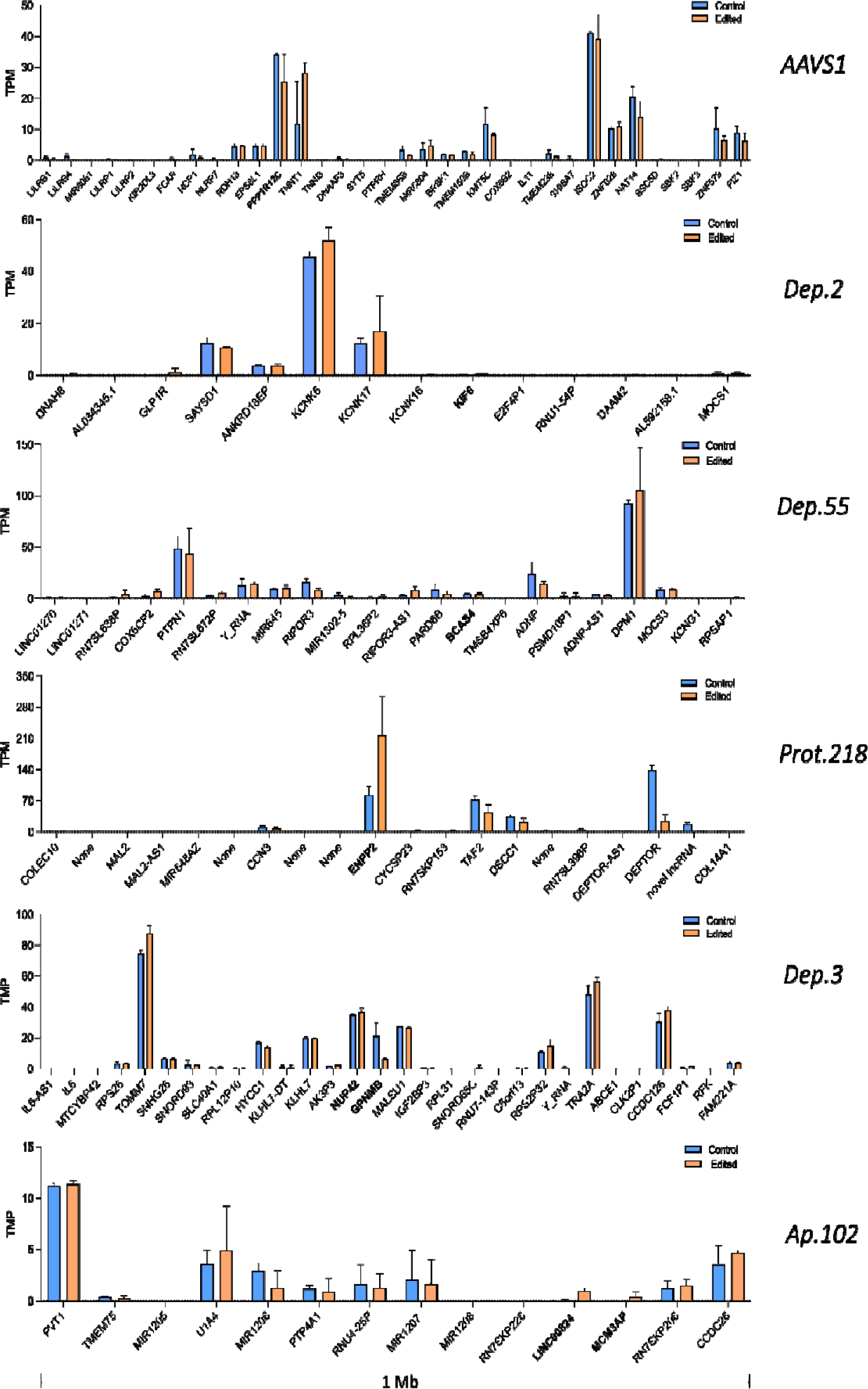
Gene expression changes within 1 Mb surrounding the insertion sites. We used normalized expression values (transcript per million, TPM) from RNAseq to depict transcriptional changes within 1 Mb around the insertion sites. For intronic GSH sites the host gene is bold. For intergenic GSH sites the flanking genes are bold. The comparison before (blue) and after (orange) gene addition displayed minimal or no disruptions of transcriptional units surrounding *AAVS1*, *Dep.2*, *Dep.55*, *Prot.218*, *Dep.3* and *Ap108* GSH sites. In the *Proto.218* locus a fair up-regulation of the host gene (*ENPP2*) was detected. Mean ± s.d. *n* = 2 independent human donors.

**Fig. S4.**
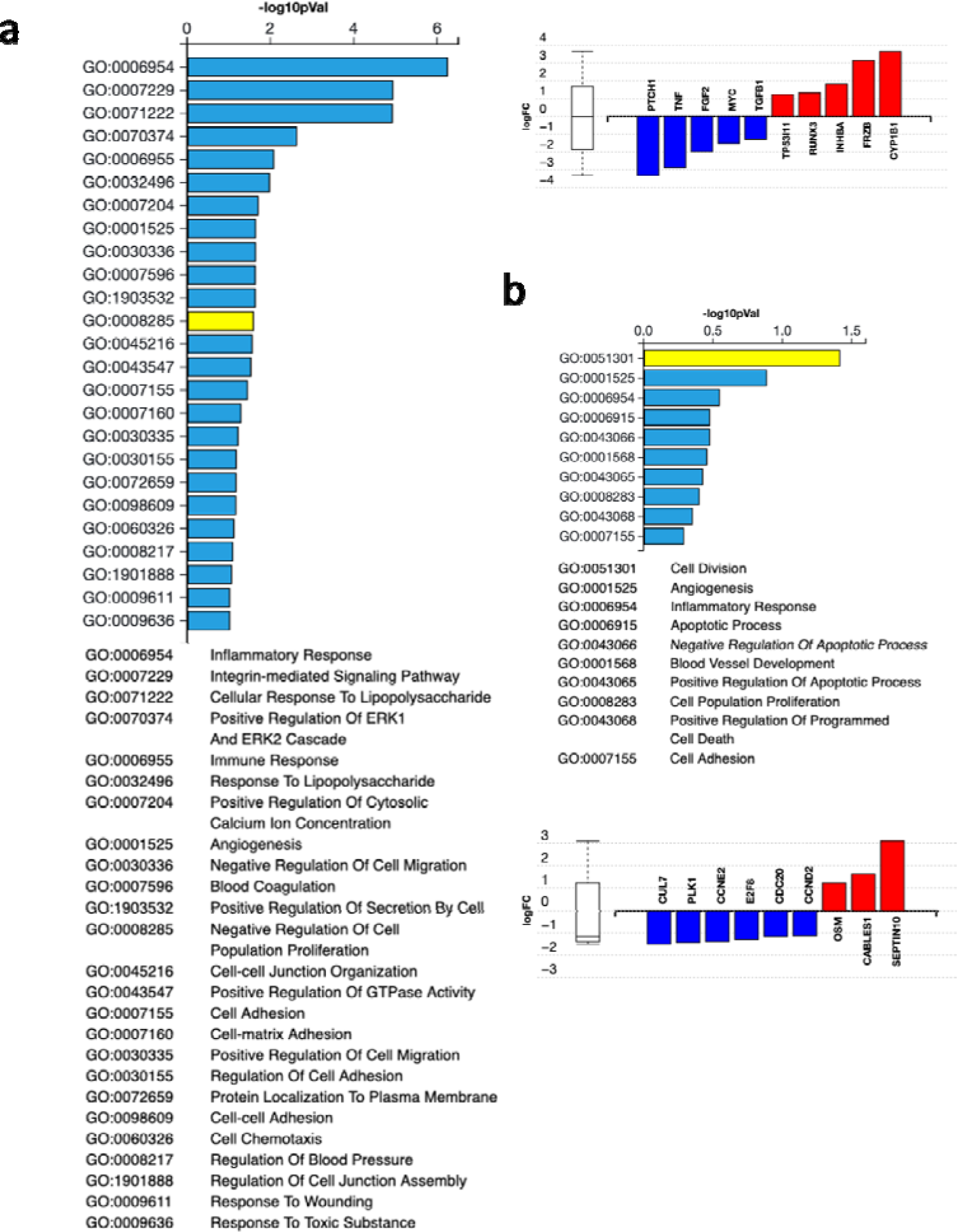
Gene ontology annotations for *AAVS1*- and *Dep.55*-targeted cells. **A** *AAVS1*. **b** *Dep.55*. Both GSH-edited cells include annotated genes to regulate cell proliferation (yellow), however, for Dep.55 GSH site those genes represent the first featured group suggesting tight regulation of cell proliferation in vitro.

**Fig. S5.**
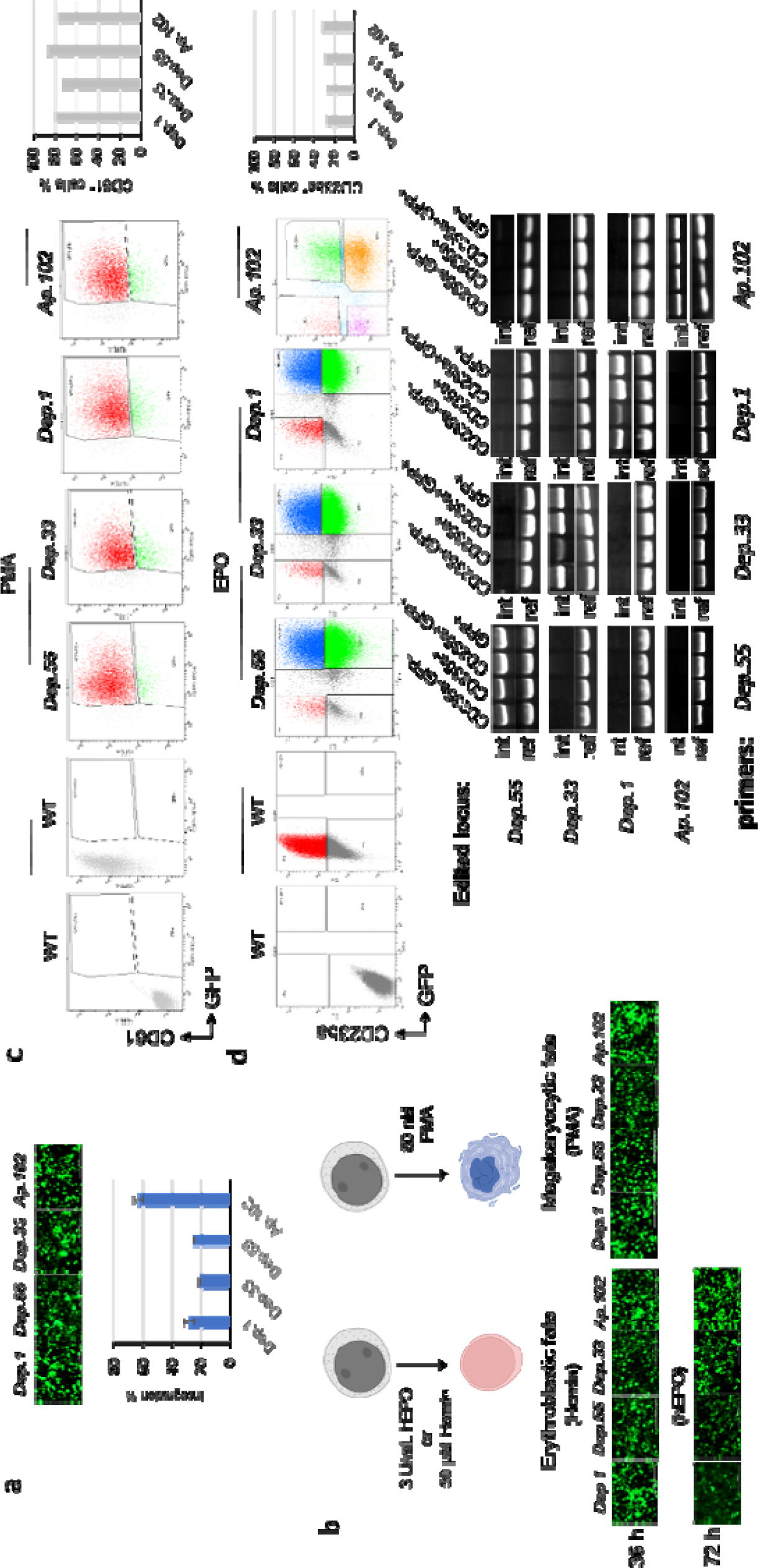
Erythroid and megakaryocytic induction of *Dep.1*, *Dep.55*, *Ap.102*, or *Dep.33* knocked in K562 cells. **a.** Stable cells were generated through delivery of ssDNA templates and the appropriated CRIPSR:gRNA complexes. The percentage of edited alleles was determined through ddPCR similarly to CD34^+^ HSPCs (Supplementary Table 10). Given that K562 is triploid, a 30% of integration indicates mostly monoallelic integration. **b** Modified K562 cells were induced to erythroid or megakaryocytic phenotypes by stimulus with 3 U/mL hEPO (or 50 μM Hemin) and 80 nM PMA, respectively. Representative pictures of induced cells at indicated times are showed. **c** Cell sorting detected up to 90% of CD61+ megakaryocytic cells with sustained expression of *eGFP* (representative dot plots and corresponding bar plot are showed). **d** Erythroid induction reached 30% CD235a^+^ cells after 3 days of hEPO treatment. Sorted phenotypes (i.e., CD235a^-^GFP^-^, CD235a^+^, CD235a^+^GFP^+^, and GFP^+^) were processed through PCR to confirm the presence of the transgene and discard effects produced by potentially random integrated clones (below). As shown the transgene was detected according to the stably edited K562 cell line (i.e. *Dep.1*, *Dep.33*, *Dep.55* or *Ap.102*), whereas the reference amplicon was observed independently of the analyzed sample. “int” refers to integration primers, and “ref” corresponds to reference primers which sequences are available in Supplementary Table 7).

**Fig. S6.**
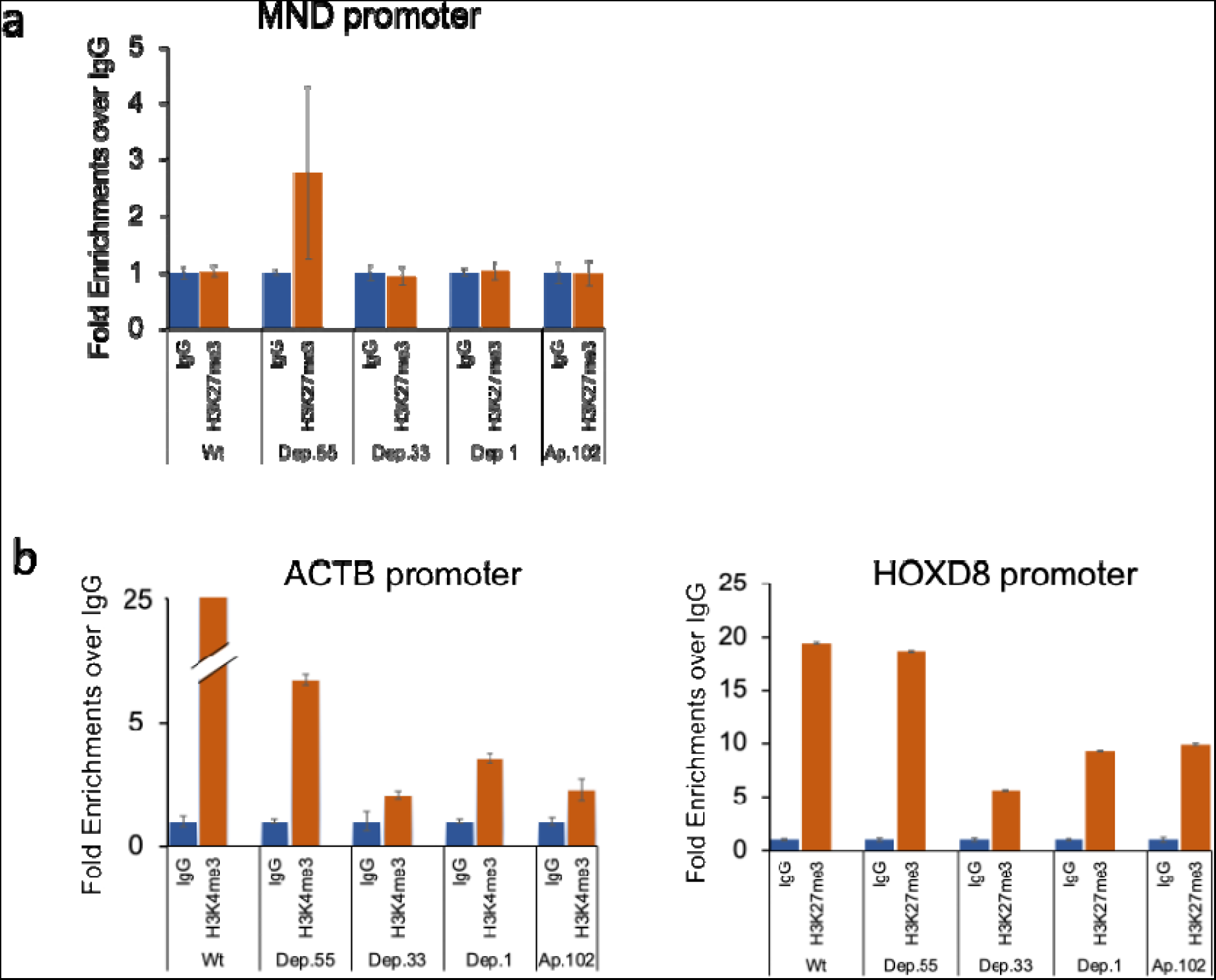
H3K27me3 shows no enrichment at the transgene in induced megakaryocytic cells a.) Enrichment of H3K27me3 at the MND transgene promoter. b) Positive control enrichments of H3K4me3 at the *ACTB* locus and H3K27me3 at the HoxD8 locus

