## Supplemental Table 1 for "Development of evolutionarily conserved viral integration sites as safe harbors for human gene therapy" (from another version of this preprint: version 2, 2023-09-25)

| EVE | Type | Chromosome | Gene1 | Gene2 |
|---|---|---|---|---|
| Control-PPP1R12C-AAVS1 | Control | chr19 | PPP1R12C | PPP1R12C |
| dependo.0-macropus | Intergenic | chr16 | CDH8 | CDH11 |
| dependo.1-whippomorpha | Intronic | chr9 | PAX5 | PAX5 |
| dependo.2-vespertilionidae | Intronic | chr6 | KIF6 | KIF6 |
| dependo.3-lagomorpha | Intergenic | chr7 | NUPL2 | GPNMB |
| dependo.4-rhinocerotidae | Intergenic | chr2 | CCDC85A | VRK2 |
| dependo.5-rhinocerotidae | Intronic | chr4 | USP38 | USP38 |
| dependo.6-elephas | Intergenic | chr18 | PIK3C3 | CELF4 |
| dependo.7-bradypus? | | | | |
| dependo.8-dasypus | Intergenic | chr22 | PRR14L | KCNH8 |
| dependo.9-hyracoidea | | | | |
| dependo.10-diprotodontia | Intergenic | chr8 | MRPS28 | TPD52 |
| dependo.12-orycteropus | | | | |
| dependo.13-cercopithecidae | Intergenic | chr21 | HUNK | MIS18A |
| dependo.14-cercopithecidae | Intronic | chr12 | KRT6A | KRT6A |
| dependo.15-colobus | Intergenic | chr3 | ZBTB20 | GAP43 |
| dependo.16-canidae | Intergenic | chr2 | SPATS2L | MAIP1 |
| dependo.20-laurasiatheria | Intronic | chr22 | GGT5 | GGT5 |
| dependo.22-laurasiatheria | Intergenic | chrX | DIAPH2 | PCDH19 |
| dependo.23-camelidae | Intergenic | chr11 | PDGFD | CASP12 |
| dependo.26-strepsirrhini | Intergenic | chr11 | ENSPSMG000 | ACAD8 |
| dependo.27-eulemur | Intergenic | chr5 | PRR16 | SRFBP1 |
| dependo.28-daubentonia | Intergenic | chr12 | LRRK2 | CNTN1 |
| dependo.33-pteropodidae | Intergenic | chr5 | PARP8 | ISL1 |
| dependo.34-megadermatvespertilion | Intergenic | chr3 | RPL18AP8 | RCC2P5 |
| dependo.35-vespertilionidae | Intronic | chr7 | ANKRD7 | ANKRD7 |
| dependo.36-phyllostomidae | Intergenic | chr6 | TMEM200A | SMLR1 |
| dependo.42-chinchilla | Intergenic | chr17 | PLEKHM1 | LRRC37A4P |
| dependo.43-octodon | Intergenic | chr22 | APOL1 | MYH9 |
| dependo.45-bathyergidae | Intergenic | chr6 | SLC18B1 | EYA4 |
| dependo.46-gliridae | Intronic | chr9 | PCSK5 | PCSK5 |
| dependo.47-pedetes | Intergenic | chr9 | TLE4 | TLE1 |
| dependo.48-mus | Intergenic | chr16 | CES1P2 | CES1P1 |
| dependo.49-apodemus | Intronic | chr9 | PCSK5 | PCSK5 |
| dependo.50-muroidae | Intergenic | chr16 | SLC6A2 | CES1P1 |
| dependo.51-muridae | Intergenic | chr8 | SNX16 | RALYL |
| dependo.54-cavid | Intergenic | chr22 | APOL3 | APOL4 |
| dependo.55-rodent | Intronic | chr20 | BCAS4 | BCAS4 |
| dependo.56-cavid | Intronic | chr3 | GUCA1C | GUCA1C |
| dependo.58-castor | Intergenic | chr2 | LRRTM4 | REG3G |

| | | | | |
|---|---|---|---|---|
| dependo.61-muscardinus | Intergenic | chr2 | CDH18 | CDH12 |
| dependo.160-gymnobelideus | Intergenic | chr9 | SEC61B | NAMA |
| dependo.190-sarcophilus | Intronic | chr3 | SGO1-AS1 | SGO1-AS1 |
| dependo.201-thamnophis | Intergenic | chr18 | GALR1 | SALL3 |
| dependo.221-pelusios | Intronic | chr1 | PIAS3 | PIAS3 |
| dependo.365-passeriformes | Intergenic | chr10 | ANKRD1 | PCGF5 |
| dependo.366-passeriformes | Intergenic | chr4 | PTPN13 | AFF1 |
| dependo.367-otididae | Intergenic | chr12 | TMTC2 | SLC6A15 |
| dependo.370-pelecanus | Intergenic | chr10 | KLF6 | AKR1E12 |
| dependo.371-egretta | Intergenic | chr12 | PDE3A | AEBP2 |
| dependo.372-opisthocomus | Intergenic | chr6 | SLC35F1 | CEP85L |
| dependo.xx-scolopacidae | Intergenic | chr12 | KRAS | LMNTD1 |
| dependo.57-ornithorhynchus | Intergenic | chr2 | MFSD9 | TGFBRAP1 |
| dependo.59-ornithorhynchus | Intergenic | chr6 | CITED2 | NMBR |
| dependo.61-sarcophilus | Intronic | chr3 | MECOM | MECOM |
| dependo.62-sarcophilus | Intergenic | chr3 | TUSC7 | IGSF11 |
| dependo.xx-mus | Intergenic | chr1 | SERTAD4 | HHAT |
| amdo.1-ellobius | Intergenic | chr3 | IGF2BP2 | TRA2B |
| amdo.2-ellobius | Intronic | chr17 | MAFG | MAFG |
| ap.101-serpentes-UR | Intergenic | chr12 | R3HDM2 | INHBC |
| ap.105-ptyas | Intronic | chr11 | GRM5 | GRM5 |
| ap.102-procavia | Intergenic | chr8 | PVT1 | CCDC26 |
| erythyro.2-dolichotis | Intergenic | chr7 | AKNRD7 | KCND2 |
| erythyro.1-indri | Intronic | chrX | IL1RAPL1 | IL1RAPL1 |
| proto.2-MusSpr | Intronic | chr12 | PDZRN4 | PDZRN4 |
| proto.4-MusSpi | Intronic | chr7 | CCDC146 | CCDC146 |
| proto.xx-marsupials | Intronic | chr8 | ENPP2 | ENPP2 |
| proto.1-RatNor | Intergenic | chr9 | CAVIN4 | PLPPR1 |
| proto.101-MonDom | Intergenic | chr10 | ADARB2 | PFKP |
| proto.103-MonDom | Intergenic | chr9 | LINGO2 | ACO1 |
| proto.104-MonDom | Intergenic | chr10 | ADARB2 | PFKP |
| proto.101-104-MonDom | Intergenic | chr10 | ADARB2 | PFKP |
| proto.131-Ctenomyidae | Intergenic | chr1 | CDC73 | KCNT2 |
| proto.132-Ctenomyidae | Intronic | chr1 | COL24A1 | COL24A1 |
| proto.151-diprodontia | Intronic | chr13 | KLHL1 | KLHL1 |
| proto.152-Sarcophilus | Intergenic | chr5 | MRPL22 | SGCD |
| proto.154-Phascolarctos | Intronic | chr21 | KCNJ6 | KCNJ6 |

| | | | | |
|---|---|---|---|---|
| proto.158-Macropus | Intronic | chr11 | ME3 | ME3 |
| proto.161-MacEug | Intronic | chr17 | CA10 | CA10 |
| proto.163-Phascolarctos | Intronic | chr17 | CA10 | CA10 |
| proto.yy-marsupials | Intronic | chr17 | CA10 | CA10 |
| proto.164-Macropus | Intergenic | chr8 | AGPAT5 | XKR5 |
| proto.167-Vombatus_ursinus | Intronic | chr10 | MANCR | MANCR |
| proto.172-MonDom | Intergenic | chr6 | TBX18 | NT5E |
| proto.174-OctDeg | Intergenic | chr9 | PHF2 | BARX1 |
| proto.176-NanGal | Intronic | chr7 | POT1-AS1 | POT1-AS1 |
| proto.180-PhaCin | Intronic | chr8 | ENPP2 | ENPP2 |
| proto.bb-marsupials | Intronic | chr10 | MANCR | MANCR |
| proto.181-PhaCin | Intronic | chr10 | MANCR | LINC00705 |
| proto.182-PhaCin | Intronic | chr15 | BTBD1 | BTBD1 |
| proto.183-PhaCin | Intronic | chr3 | VPS8 | VPS8 |
| proto.184-PhaCin | Intronic | chr9 | AGTPBP1 | AGTPBP1 |
| proto.185-PhaCin | Intergenic | chr6 | CD109 | COL12A1 |
| proto.186-PhaCin | Intergenic | chr2 | DLX2 | ITGA5 |
| proto.187-PhaCin | Intergenic | chr4 | SFRP2 | ENSG000002[illegible] |
| proto.188-PhaCin | Intergenic | chr6 | TBX18 | NT5E |
| proto.aa-marsupials | intergenic | chr6 | TBX18 | NT5E |
| proto.193-CteSoc | Intergenic | chr1 | CDC73 | KCNT2 |
| proto.194-SarHar | Intergenic | chr4 | PCDH10 | PCDH18 |
| proto.196-SarHar | Intergenic | chr5 | DAB2 | PTGER4 |
| proto.201-ChrAsi | Intronic | chr7 | CCDC146 | CCDC146 |
| proto.202-ChrAsi | Intronic | chr15 | CEMIP | CEMIP |
| proto.209-MacEug | Intronic | chr3 | VPS8 | VPS8 |
| proto.zz-marsupials | Intronic | chr3 | VPS8 | VPS8 |
| proto.215-MacEug | Intronic | chr3 | ZPLD1 | ZPLD1 |
| proto.217-VomUrs | Intergenic | chr6 | TBX18 | NT5E |
| proto.218-VomUrs | Intronic | chr8 | ENPP2 | ENPP2 |

| HumanCoords | MinAge | VirusGenus |
|---|---|---|
| NA | NA | NA |
| 16:62189064-62190580 | 0 | Dependoparvovirus |
| 9:36853489-36853496 | 53.75 | Dependoparvovirus |
| 6:39422982-39425107 | 31.32 | Dependoparvovirus |
| 7:23218501-23220555 | 51.43 | Dependoparvovirus |
| 2:56547287-56548087 | 14.12 | Dependoparvovirus |
| 4:143184917-143224429 | 14.12 | Dependoparvovirus |
| 18:38855055-38866235 | 0 | Dependoparvovirus |
| NA | | Dependoparvovirus |
| 22:19148510-31750132 | 0 | Dependoparvovirus |
| NA | | Dependoparvovirus |
| 8:80030259-80034745 | 35.01 | Dependoparvovirus |
| NA | | Dependoparvovirus |
| 21:32098068-32098075 | 12.4 | Dependoparvovirus |
| 21:52482109-52482121 | 12.4 | Dependoparvovirus |
| 3:115526836-115532441 | 0 | Dependoparvovirus |
| 2:200143437-200147284 | 14.15 | Dependoparvovirus |
| 22:24236476-24242627 | 77.75 | Dependoparvovirus |
| X:98175245-98175812 | 77.75 | Dependoparvovirus |
| 11:104771681-104771952 | 20.56 | Dependoparvovirus |
| 11:134272095-134345651 | 54.69 | Dependoparvovirus |
| 5:120916921-120917619 | | Dependoparvovirus |
| 12:40658392-40660499 | | Dependoparvovirus |
| 5:50958291-50958438 | 33.76 | Dependoparvovirus |
| 3:96720524-96720525 | 61.66 | Dependoparvovirus |
| 7:118263040-118263054 | 33.24 | Dependoparvovirus |
| 6:130764851-130765531 | | Dependoparvovirus |
| 17:45490775-45505883 | 0 | Dependoparvovirus |
| 22:36268100-36281236 | 0 | Dependoparvovirus |
| 6:133180282-133183376 | 43.36 | Dependoparvovirus |
| 9:76003270-76011839 | NA | Dependoparvovirus |
| 9:80967898-80971940 | NA | Dependoparvovirus |
| 16:55726519-55793611 | NA | Dependoparvovirus |
| 9:76005001-76010043 | NA | Dependoparvovirus |
| 16:55706192-55760599 | NA | Dependoparvovirus |
| 8:83010298-83038101 | NA | Dependoparvovirus |
| 22:36134752-36197047 | 43.36 | Dependoparvovirus |
| 20:50800440-50801000 | 43.36 | Dependoparvovirus |
| 3:108913357-108913358 | 5.66 | Dependoparvovirus |
| 2:78044772-78060998 | 0 | Dependoparvovirus |

| | | |
|---|---|---|
| 5:20680471-20684922? | NA | Dependoparvovirus |
| 9:99239314-99292227 | 0 | Dependoparvovirus |
| 3:20858652-20870649 | 0 | Dependoparvovirus |
| 18:78777575-78805227 | | Dependoparvovirus |
| 1:145857026-145867329 | 0 | Dependoparvovirus |
| 10:90998918-91163307 | 44 | Dependoparvovirus |
| 4:86815140-86959933 | 65.69 | Dependoparvovirus |
| 12:84666026-84860662 | | Dependoparvovirus |
| 10:4512911-4526527 | 0 | Dependoparvovirus |
| 12:19520263-19685396 | 0 | Dependoparvovirus |
| 6:118317676-118460772 | 0 | Dependoparvovirus |
| 12:25250929-25648604 | 39 | Dependoparvovirus |
| 2:103605381-103618136 | 0 | Dependoparvovirus |
| 6:139599746-139606611 | 0 | Dependoparvovirus |
| 3:169514067-169525076 | 0 | Dependoparvovirus |
| 3:118743559-118744184 | 0 | Dependoparvovirus |
| 1:182539754-182540179 | 0 | Dependoparvovirus |
| 3:185825042-185914558 | 0 | Amdoparvovirus |
| 17:81918270-81927735 | 0 | Amdoparvovirus |
| 12:57411074-57449700 | 29.51 | Amdoparvovirus |
| 11:88504576-89065945 | 0 | Amdoparvovirus |
| 8:128435359-128437048 | 10.17 | Amdoparvovirus |
| 7:118242732-120273175 | 0 | Erythroparvovirus |
| X:29148419-29154761 | 0 | Erythroparvovirus |
| 12:41262894-41263054 | 0 | Protoparvovirus |
| 7:77247600-77252777 | | Protoparvovirus |
| 8:119591710-119593739 | NA | Protoparvovirus |
| 9:100865872-100872934 | 0 | Protoparvovirus |
| 10:1897420-1929958 | 0 | Protoparvovirus |
| 9:30918799-31106170 | 0 | Protoparvovirus |
| 10:1897420-1929958 | 0 | Protoparvovirus |
| 10:1897420-1929958 | 0 | Protoparvovirus |
| 1:194737226-194740522 | NA | Protoparvovirus |
| 1:85954973-85954090 | NA | Protoparvovirus |
| 13:69701702-70107699 | NA | Protoparvovirus |
| 5:154969411-156326757 | 0 | Protoparvovirus |
| 21:37829378-37840593 | 0 | Protoparvovirus |

| | | | |
|---|---|---|---|
| 11:86493907-86497959 | | 0 | Protoparvovirus |
| 17:51847456-51858255 | | 0 | Protoparvovirus |
| 17:51847419-51851769 | | 0 | Protoparvovirus |
| 17:51847419-51851769 | NA | | Protoparvovirus |
| 8:6759763-6808517 | | 0 | Protoparvovirus |
| 10:4652078-4656006 | | 0 | Protoparvovirus |
| 6:85396039-85450172 | | 0 | Protoparvovirus |
| 9:93871473-93890849 | | 0 | Protoparvovirus |
| 7:125056826-125056864 | | 0 | Protoparvovirus |
| 8:119591710-119593739 | | 0 | Protoparvovirus |
| 10:4652078-4656006 | NA | | Protoparvovirus |
| 10:4652078-4656002 | | 0 | Protoparvovirus |
| 15:83067252-83016423 | | 0 | Protoparvovirus |
| 3:184812143-184915483 | | 0 | Protoparvovirus |
| 9:85548005-85575207 | | 0 | Protoparvovirus |
| 6:74854532-74870040 | | 0 | Protoparvovirus |
| 2:172298255-172298513 | | 0 | Protoparvovirus |
| 4:153851339-153886501 | | 0 | Protoparvovirus |
| 6:85345929-85450148 | | 0 | Protoparvovirus |
| 6:85345929-85450148 | | 0 | Protoparvovirus |
| 1:194737226-194740522 | | 0 | Protoparvovirus |
| 4:133194700-137518918 | | 0 | Protoparvovirus |
| 5:39424980-40679915 | | 0 | Protoparvovirus |
| 7:77241899-77241990 | | 0 | Protoparvovirus |
| 15:80870154-80870954 | | 0 | Protoparvovirus |
| 3:184915484-184918417 | | 0 | Protoparvovirus |
| 3:184812143-184915483 | NA | | Protoparvovirus |
| 3:102394557-102404994 | | 0 | Protoparvovirus |
| 6:85514953-85355198 | | 0 | Protoparvovirus |
| 8:119591710-119593739 | | 0 | Protoparvovirus |
